## Supplementary Material for "Omissions of Threat Trigger Subjective Relief and Prediction Error-Like Signaling in the Human Reward and Salience Systems"

### Supplementary Materials

#### Supplementary Methods

##### Participant sample characteristics

**Supplementary Note 1: Sample size rationale**

The study was the first fMRI application of the EVA task. As such, we were not able to conduct a power analyses based on previously reported fMRI effects. Therefore, to ensure enough statistical power to our analysis, we recruited $n$ = 30 healthy volunteers to participate in our study, which is a commonly used sample size for fMRI research. This sample size furthermore exceeds the required sample size for replicating the behavioral findings based on a power of .95 and alpha of .05, and effect sizes reported in Willems & Vervliet, 2021. Specifically, effect sizes for the main effects of Probability (within the Probability x Block RM-ANOVA) and Intensity (within Probability x Intensity x Block RM-ANOVA) in this validation study were $\eta_p^2$ = .62, $\varepsilon$ = 0.52 and $\eta_p^2$ = .81, $\varepsilon$ = 0.62 for relief-pleasantness ratings, and $\eta_p^2$ = .72 and $\eta_p^2$ = .48, $\varepsilon$ = 0.67 for omission-induced SCR, respectively. Our calculations (G*power software, selecting F tests, ANOVA: repeated measures, within factors, for 4 levels of probability and 3 levels of intensity) revealed that for relief-pleasantness, a sample size of $n$ = 9, and for omission SCR a sample size of $n$ = 14 was sufficient to replicate the probability and intensity effects we observed in the subjective and physiological data with a power of .95 and an alpha of .05.

**Supplementary Note 2: Exclusion Criteria**

Only healthy, right-handed individuals between the ages of 16 and 30, and with a body-mass index of 18.5-25 kg/m² were eligible to participate in the study. Female participants were furthermore required to take hormonal contraceptives or to be tested in the follicular phase of their menstrual cycle.

In addition, participants were screened upon enrollment for the following exclusion criteria:

- Past/present medical disorders: neurological, cardiovascular or respiratory disorders, hypertension, migraine, head trauma with loss of consciousness, chronic (duration of > 3 months) or acute pain;
- Past/present psychiatric disorders: clinical depression, anxiety, psychotic disorders, mood disorders, eating disorders, somatoform disorders, substance-related disorders or any other psychiatric disorder;

- Regular medication use (except from oral contraceptives): Treatment in the last 6 months with antidepressants, antipsychotics, sedative hypnotics, psychostimulants, glucocorticoids, appetite suppressants, estrogens, opiates (such as pain medication and coughing syrup with codeine), or dopaminergic medications;
- Being pregnant or lactating;
- Request from medical doctor to stay away from stressful situations;
- Smoking;
- Cannabis use or any other drug of abuse (in the last 12 months);
- Regular high alcohol (> 10 units/week), caffeine (>1000 ml coffee daily or equivalent) or energy drink (> 1 drink/day) intake
- Any contraindication for MRI: e.g., cochlear implant, cardiac pacemaker, neural stimulator, metallic body inclusion or any other metal (implanted) in the body which may interfere with MRI scanning; claustrophobia or severe back problems that may interfere with complying to scanning procedures.

Demographic information related to the final sample can be found in Supplementary Table 1 and 2

**Supplementary Table 1: Demographics of Included Participants**

| Subject ID | Age | Gender | Excluded from rating analysis | Excluded from SCR analysis | Excluded from fMRI analysis |
|---|---|---|---|---|---|
| Sub-01 | 24 | Male | | | |
| Sub-02 | 19 | Female | | | |
| Sub-03 | 21 | Female | | | Run 1 |
| Sub-04 | 25 | Female | | | |
| Sub-05 | 19 | Female | | | |
| Sub-06 | 20 | Male | | | |
| Sub-07 | 19 | Female | | | |
| Sub-08 | 24 | Female | | | |
| Sub-09 | 22 | Male | | | |
| Sub-10 | 21 | Female | | All runs (non-responder) | |
| Sub-13 | 20 | Male | | All runs (non-responder) | |
| Sub-14 | 20 | Male | Run 4 | Run 4 | Run 4 |
| Sub-15 | 19 | Female | | First 4 trials of Run 1 | |
| Sub-16 | 18 | Male | | | |
| Sub-20 | 25 | Female | | | |
| Sub-21 | 19 | Male | Run 4 | All runs (technical difficulties) | Run 4 |
| Sub-22 | 20 | Female | | All runs (non-responder) | |
| Sub-23 | 23 | Female | | | |
| Sub-24 | 25 | Male | | | |
| Sub-25 | 25 | Male | | All runs (non-responder) | |
| Sub-26 | 25 | Male | | | |
| Sub-27 | 22 | Female | | | |
| Sub-29 | 19 | Female | | | |
| Sub-30 | 19 | Female | | | |
| Sub-31 | 18 | Female | | | |
| Sub-32 | 18 | Male | | | |
| Sub-33 | 19 | Female | Last trial of Run 4 | Last trial of Run 4 | Run 4 |
| Sub-34 | 18 | Female | | | |
| Sub-35 | 18 | Male | | | |
| Sub-36 | 18 | Female | | | |
| Sub-37 | 18 | Female | | | |
| **TOTAL (N = 31)** | ***M* = 20.65** | **19 Females** | **Final sample: N = 31** | **Final sample: N = 26 (17 females)** | **Final sample: N = 31** |

**Supplementary Table 2: Descriptives of Questionnaire scores**

| **Questionnaire scale** | **Mean** | **Standard deviation** | ***N*** |
|---|---|---|---|
| STAI Trait | 36.68 | 9.25 | 31 |
| IUS | 63.20 | 13.46 | 30 |
| Prospective subscale | 17.47 | 4.95 | 30 |
| Inhibitory subscale | 10.20 | 3.36 | 30 |
| DTS | 3.69 | 0.67 | 31 |
| Absorption subscale | 3.59 | 0.87 | 31 |
| Appraisal subscale | 3.85 | 0.77 | 31 |
| Regulation subscale | 3.46 | 0.79 | 31 |
| Tolerance subscale | 3.87 | 0.68 | 31 |
| DASS | | | |
| Depression subscale | 7.00 | 6.23 | 30 |
| Anxiety subscale | 2.97 | 3.58 | 30 |
| Stress subscale | 7.43 | 5.38 | 30 |
| ASI | 14.97 | 8.16 | 30 |
| Physical subscale | 4.10 | 3.40 | 30 |
| Cognitive subscale | 3.23 | 3.45 | 30 |
| Social subscale | 7.63 | 4.07 | 30 |
| LOTR | | | |
| Optimism subscale | 7.10 | 2.49 | 31 |
| Pessimism subscale | 5.19 | 2.51 | 31 |
| BAS | | | |
| Drive subscale | 11.45 | 2.03 | 31 |
| Fun seeking subscale | 12.03 | 2.01 | 31 |
| Reward responsiveness subscale | 17.26 | 1.95 | 31 |
| BIS | 20.71 | 3.73 | 31 |
| PANAS | | | |
| Positive subscale | 37.10 | 3.84 | 31 |
| Negative subscale | 21.71 | 5.72 | 31 |
| PCS | | | |
| Helplessness subscale | 4.50 | 2.96 | 30 |
| Magnification subscale | 2.90 | 1.97 | 30 |
| Rumination subscale | 7.73 | 2.78 | 30 |

*Note.* Participants filled out a questionnaire battery during the intake session, as part of a larger attempt to relate individual differences in anxiety- and pain-related traits to individual differences in relief. The battery consisted of Dutch versions of the Depression, Anxiety & Stress Scales (DASS)(De Beurs & Van Dyck, 2001; Lovibond & Lovibond, 1995), State-Trait Anxiety Inventory (STAI)(Spielberger et al., 1983; Van der Ploeg, 1982), Intolerance of Uncertainty Scale (IUS)(de Bruin et al., 2006; Freeston et al., 1994), Positive and negative Affect Schedule (PANAS)(Engelen et al., 2006; Watson et al., 1988), Behavioral Inhibition Scale and Behavioral Activation Scale (BIS/BAS)(Franken et al., 2005; White & Carver, 1994), Distress Tolerance Scale (DTS)(Simons & Gaher, 2005) and Life Optimism Trait – Revised (LOT-R)(Klooster et al., 2010; Scheier et al., 1994), the Anxiety Sensitivity Index (ASI)(Taylor et al., 2007), and the Pain Catastrophizing Scale (PCS)(Sullivan et al., 1995). *N* represents number of participants that completed the questionnaire and that was used to calculate the summary statistics.

**Experimental Task**

**Supplementary Note 3: Stimulation Workup Procedure**

Participants were presented with a range of increasingly intense stimulations and were asked to rate their unpleasantness on a scale from 0 (no sensation) to 10 (extreme, intolerable pain), with a rating of 1 corresponding to "clear sensation", a rating of 3 to a "mildly uncomfortable sensation", a rating of 5 to a "very uncomfortable, but not painful sensation", a rating of 6 to "faint pain", a rating of 7 to "pain" and a rating of 8 to "significant, but tolerable pain". While participants were asked to select a stimulus that was significantly painful, but tolerable (rated as an 8), the researcher additionally selected two other intensities corresponding to a rating of 3 and 5.

**Supplementary Figure 1: Trial Types and Numbers**

| | Intensity | | | | | | | |
|---|---|---|---|---|---|---|---|---|
| | Weak | | Moderate | | strong | | | |
| Probability | Omission | Stimulation | Omission | Stimulation | Omission | stimulation | | |
| 0% | | | *No intensity info* | | | | 12 | 0 |
| 25% | 4 | 1 (+1*) | 4 | 1 | 4 | 1 | 12 | 4 |
| 50% | 4 | 1 | 4 | 1 (+1*) | 4 | 1 | 12 | 4 |
| 75% | 4 | 1 | 4 | 1 | 4 | 1 (+1*) | 12 | 4 |
| 100% | 0 | 4 | 0 | 4 | 0 | 4 | 0 | 12 |
| Total | 12 | 8 | 12 | 8 | 12 | 8 | 48 | 24 |

** This is an example. In general there should be one additional weak, one additional moderate, and one additional strong stimulation with exactly one having probability 25%, one having probability 50%, and one having probability 75%*

**Supplementary Note 4: "Accurate" probability instructions do not alter the Probability-effect**

A question that was raised by the reviewers was whether the inconsistency between the probability instruction and the experienced reinforcement rate could have detrimental effects on the Probability-related results; especially because the effect of Probability was smaller when only including non-0% trials.

However, there are good reasons to believe that the relatively smaller difference between 25% to 75% trials was not caused by the “inaccurate” nature of our instructions, but that they are inherent to “uncertain” probabilities.

First, in a previously unpublished pilot study, we provided participants with “accurate” probability instructions, meaning that the instruction corresponded to the actual reinforcement rate (e.g., 75% instructions were followed by a stimulation in 75% of the trials etc.). In line with the present results and our previous behavioral study (Willems & Vervliet, 2021), the results of this pilot (N = 20) showed that the difference in the reported relief between the different probability levels was largest when comparing 0% and the rest (25%, 50% and 75%). Furthermore the overall effect size of Probability (excluding 0%) matched the one of our previous behavioral study (Willems & Vervliet, 2021): ηp2 = +/- 0.50.

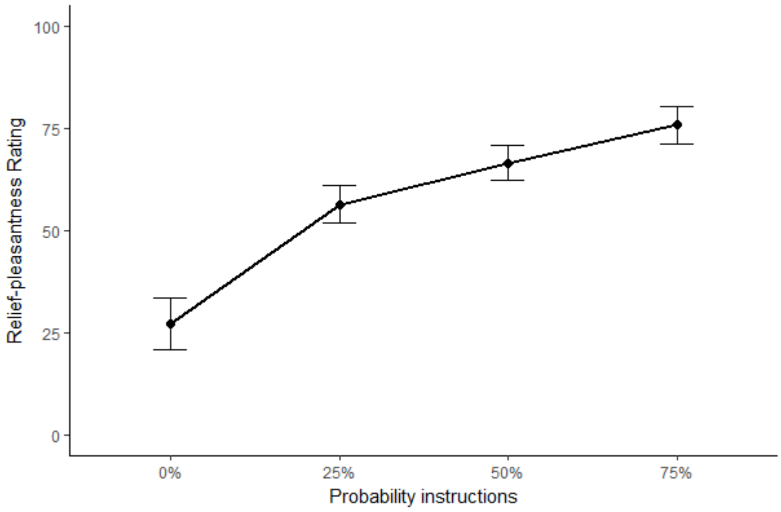


Main effect of Probability including 0% : F(1.74,31.23) = 53.94, p < .001, ηp2 = 0.75

Main effect of Probability excluding 0%: F(1.50, 28.43) = 21.03, p < .001, ηp2 = 0.53

Second, also in other published studies that used CSs with varying reinforcement rates (which either included explicit written instructions of the reinforcement rates or not) showed that the difference in expectations, anticipatory SCR or omission SCR was largest when comparing the CS0% to the other CSs of varying reinforcement rates (Grings & Sukoneck, 1971; Öhman et al., 1973; Ojala et al., 2022).

Together, this suggests that when there is a possibility of stimulation, any additional difference in probability will have a smaller effect on the omission responses, irrespective of whether the underlying reinforcement rate is accurate or not.

**Statistical Analyses**

**Supplementary Note 5: Deviations from pre-registration**

Main analyses for this study were pre-registered on Open Science Framemwork (OSF) and made available online (https://osf.io/ugkzf). While the general analysis approach was maintained, a number of small deviations from the pre-registered plan were made in the paper. Here, we provide an overview of these deviations.

- In addition to the predictors-of-interest we specified in the pre-registration (Probability and Intensity), we entered Run (four levels: 1, 2, 3, 4) (only in rating and SCR models) and average stimulation-unpleasantness (centered) as predictors of no-interest, as these increased model fit (as assessed via AIC). Note that Run-specific intercepts were included in the first-level GLMs of the fMRI data, but the effects of Run were not explicitly tested.

- We specified in the pre-registration that we would smooth our preprocessed functional data for the univariate analysis, however we also used the smoothed data for the multivariate analyses, since previous research has shown that this does not have a detrimental effect and might even improve the results (Hendriks et al., 2017; Op de Beeck, 2010).

- We specified in the pre-registration that all regressors in the subject-level GLMs would be modelled as stick-functions (duration = 0). However, in the paper we only modeled the outcome (omission/shock) as stick regressors (similar to previous research), whereas the other regressors (instructions, ratings) were modeled as boxcar regressors to remove their variance from the implicit baseline. Furthermore, we did not include a separate regressor for the onset of the clock as it overlapped with the regressor of the instructions.

- We specified in the pre-registration that the effects of Intensity and Probability on the fMRI data would be analyzed via an full-factorial F-test in SPM12. However, to keep this analysis comparable to our rating and SCR analyses, we extracted parameter estimates from each ROI for each Probability x Intensity combination in the paper, and entered these estimates in a Linear Mixed Model including Probability, Intensity, and their interaction as regressors-of-interest, and average stimulation unpleasantness (centered) as regressor of no-interest.

- We specified in the pre-registration that relief-ratings would be entered as parametric modulator in order to assess trial-by-trial correlations between fMRI data and relief ratings. In the paper, the ratings were z-scored before entering them into the model.

- We specified in the pre-registration that we would use an 8-fold cross-validation for the LASSO-PCR. However, in the paper, we used a 5-fold cross-validation instead, in line with lab standards. Furthermore, in addition to using virtual lesions, we applied a bootstrapping approach to assess the contribution of individual regions to the model signature.

- We added the red nucleus and habenula to the secondary ROI mask, because of their close spatial proximity to the dopaminergic midbrain (red nucleus), their efferent connection to the dopaminergic midbrain (habenula) and their functional relation to omission processing in the past (Hikosaka, 2010; Linnman et al., 2011).

- We specified in the pre-registration that we would explore threat omission processing in an extended ROI mask using a voxel-wise, FDR small volume corrected approach, followed by a whole-brain parcel-wise approach. However, instead of the whole-brain parcel-wise approach, we used a voxel-wise approach within a grey matter mask, using FWE-correction. Furthermore, as for our main analyses we extracted averaged beta-estimates from each omission-processing cluster (identified with a cluster threshold of $p < .05$, FWE-corrected, following voxel threshold $p < .001$) and entered these in a Probability x Intensity LMM.

#### Supplementary Methods 1: Exploratory Rating and SCR analyses.

Besides focusing on the omission and anticipation window, we explored how Probability and Intensity instructions affected unconditioned responding (self-reported unpleasantness and stimulation SCR) to the delivery of the stimulation. We conducted two additional linear mixed models with self-reported US unpleasantness and stimulation SCR as dependent variables and Probability (2 levels: non100%, 100%) and Intensity (weak, moderate, strong), and their interaction as fixed effects, in addition to a subject-specific intercept as random effect. Note that we pooled all non100% probability levels so that the probability levels were balanced.

#### Supplementary Methods 2: Preprocessing of MRI data with fMRIPrep.

Preprocessing of the MRI data was performed using fMRIPrep 20.2.3 (Esteban, Markiewicz, et al. (2018); Esteban, Blair, et al. (2018); RRID:SCR_016216), which is based on Nipype 1.6.1 (Gorgolewski et al. (2011); Gorgolewski et al. (2018); RRID:SCR_002502).

##### *Anatomical data preprocessing.*

A total of 1 T1-weighted (T1w) images were found within the input BIDS dataset. The T1-weighted (T1w) image was corrected for intensity non-uniformity (INU) with N4BiasFieldCorrection (Tustison et al. 2010), distributed with ANTs 2.3.3 (Avants et al. 2008, RRID:SCR_004757), and used as T1w-reference throughout the workflow. The T1w-reference was then skull-stripped with a Nipype implementation of the antsBrainExtraction.sh workflow (from ANTs), using OASIS30ANTs as target template. Brain tissue segmentation of cerebrospinal fluid (CSF), white-matter (WM) and gray-matter (GM) was performed on the brain-extracted T1w using fast (FSL 5.0.9, RRID:SCR_002823, Zhang, Brady, and Smith 2001). Volume-based spatial normalization to one standard space (MNI152NLin2009cAsym) was performed through nonlinear registration with antsRegistration (ANTs 2.3.3), using brain-extracted versions of both T1w reference and the T1w template. The following template was selected for spatial normalization: ICBM 152 Nonlinear Asymmetrical template version 2009c [Fonov et al. (2009), RRID:SCR_008796; TemplateFlow ID: MNI152NLin2009cAsym],

***Functional data preprocessing.***

For each of the 4 BOLD runs found per subject (across all tasks and sessions), the following preprocessing was performed. First, a reference volume and its skull-stripped version were generated using a custom methodology of fMRIPrep. A B0-nonuniformity map (or fieldmap) was estimated based on two (or more) echo-planar imaging (EPI) references with opposing phase-encoding directions, with 3dQwarp Cox and Hyde (1997) (AFNI 20160207). Based on the estimated susceptibility distortion, a corrected EPI (echo-planar imaging) reference was calculated for a more accurate co-registration with the anatomical reference. The BOLD reference was then co-registered to the T1w reference using flirt (FSL 5.0.9, Jenkinson and Smith 2001) with the boundary-based registration (Greve and Fischl 2009) cost-function. Co-registration was configured with nine degrees of freedom to account for distortions remaining in the BOLD reference. Head-motion parameters with respect to the BOLD reference (transformation matrices, and six corresponding rotation and translation parameters) are estimated before any spatiotemporal filtering using mcflirt (FSL 5.0.9, Jenkinson et al. 2002). BOLD runs were slice-time corrected using 3dTshift from AFNI 20160207 (Cox and Hyde 1997, RRID:SCR_005927). The BOLD time-series (including slice-timing correction when applied) were resampled onto their original, native space by applying a single, composite transform to correct for head-motion and susceptibility distortions. These resampled BOLD time-series will be referred to as preprocessed BOLD in original space, or just preprocessed BOLD. The BOLD time-series were resampled into standard space, generating a preprocessed BOLD run in MNI152NLin2009cAsym space. First, a reference volume and its skull-stripped version were generated using a custom methodology of fMRIPrep. Several confounding time-series were calculated based on the preprocessed BOLD: framewise displacement (FD), DVARS and three region-wise global signals. FD was computed using two formulations following Power (absolute sum of relative motions, Power et al. (2014)) and Jenkinson (relative root mean square displacement between affines, Jenkinson et al. (2002)). FD and DVARS are calculated for each functional run, both using their implementations in Nipype (following the definitions by Power et al. 2014). The three global signals are extracted within the CSF, the WM, and the whole-brain masks. Additionally, a set of physiological regressors were extracted to allow for component-based noise correction (CompCor, Behzadi et al. 2007). Principal components are estimated after high-pass filtering the preprocessed BOLD time-series (using a discrete cosine filter with 128s cut-off) for the two CompCor variants: temporal (tCompCor) and anatomical (aCompCor). tCompCor components are then calculated from the top 2% variable voxels within the brain mask. For aCompCor, three probabilistic masks (CSF, WM and combined CSF+WM) are generated in anatomical space. The implementation differs from that of Behzadi et al. in that instead of eroding the masks by 2 pixels on BOLD space, the aCompCor masks are subtracted a mask of pixels that likely contain a volume fraction of GM. This mask is obtained by thresholding the corresponding partial volume map at 0.05, and it ensures components are not extracted from voxels containing a minimal fraction of GM. Finally, these masks are resampled into BOLD space and binarized by thresholding at 0.99 (as in the original implementation). Components are also calculated separately within the WM and CSF masks. For each CompCor decomposition, the k components with the largest singular values are retained, such that the retained components' time series are sufficient to explain 50 percent of variance across the nuisance mask (CSF, WM, combined, or temporal). The remaining components are dropped from consideration. The head-motion estimates calculated in the correction step were also placed within the corresponding confounds file. The confound time series derived from head motion estimates and global signals were expanded with the inclusion of temporal derivatives and quadratic terms for each (Satterthwaite et al. 2013). Frames that exceeded a threshold of 0.9 mm FD or 2.0 standardised DVARS were annotated as motion outliers. All resamplings can be performed with a single interpolation step by composing all the pertinent transformations (i.e. head-motion transform matrices, susceptibility distortion correction when available, and co-registrations to anatomical and output spaces). Gridded (volumetric) resamplings were performed using antsApplyTransforms (ANTs), configured with Lanczos interpolation to minimize the smoothing effects of other kernels (Lanczos 1964). Non-gridded (surface) resamplings were performed using mri_vol2surf (FreeSurfer).

Many internal operations of fMRIPrep use Nilearn 0.6.2 (Abraham et al. 2014, RRID:SCR_001362), mostly within the functional processing workflow. For more details of the pipeline, see the section corresponding to workflows in fMRIPrep's documentation.

#### Supplementary Results

##### Subjective and physiological results

**Supplementary Figure 2: Post experimental recollections of stimulation and effort to avoid future stimulations**

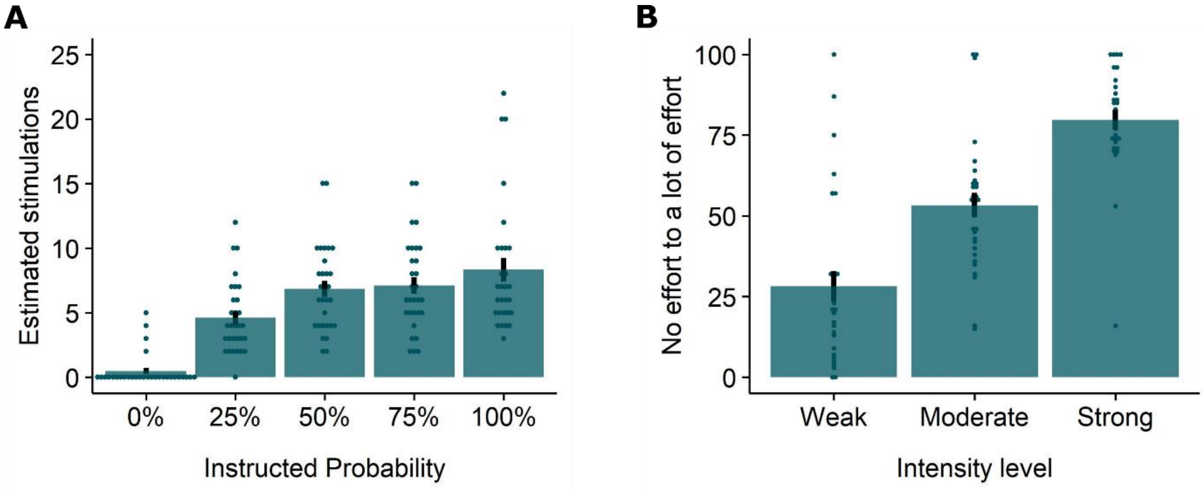


As post-experimental manipulation checks for the probability and intensity instructions, participants were asked at the end of the task how many stimuli they thought they received following instructions of each probability level (probability manipulation check), and how much effort they would exert to prevent future weak/moderate/strong stimulation (from 0 "no effort" to 100 "a lot of effort"). **A.** A Friedman test revealed that participants recollected the number of stimuli they received in line with the provided instructions ($\chi^2(4) = 69.6$, $p < .001$, W = .58). Follow-up Wilcoxon signed-rank tests indicated that participants recollected having experienced the least number of stimulations following 0% instructions (all *p'*s < .001). Furthermore, they recollected having experienced some stimulations after the 25% instructions, but less than following 50% to 100% instructions ($p < .05$). The differences between 50%, 75% and 100% did not reach significance following Bonferroni-Holm correction ($p > .13$). **B.** In line with the intensity instructions, an Intensity (weak, moderate, strong) LMM indicated that participants were willing to exert more effort to prevent stronger stimulations ($F(2, 58) = 97.22$, $p < .001$, $\omega_p^2 = 0.76$, all Bonferroni-Holm corrected pairwise comparisons, $p < .001$). Overall, these checks support that the EVA task was successful at inducing expectations of threat.

**Supplementary Figure 3: Anticipatory SCR to the instructions**

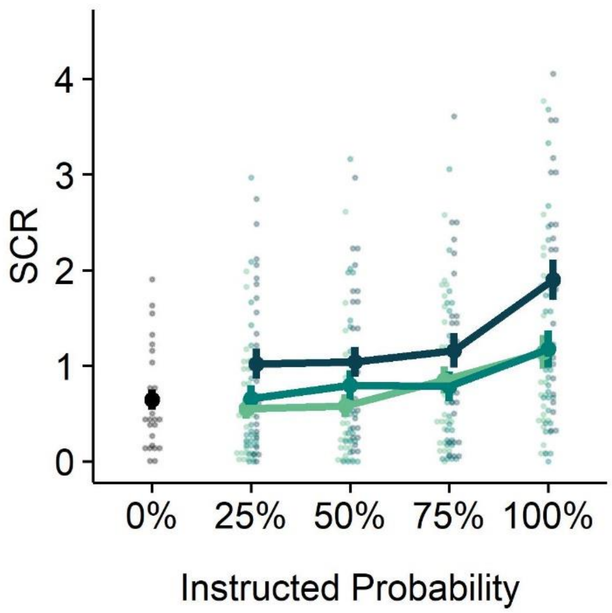


A 5 (Probability: 0%, 25%, 50%, 75%, 100%) x 4 (Run: 1, 2, 3, 4) LMM revealed that anticipatory SCR to the countdown clock increased with increasing probability instructions (main effect of Probability, $F(4, 1804) = 49.30$, $p < .001$, $\omega_p^2 = 0.10$). Bonferroni-Holm corrected follow-up pairwise contrasts confirmed that SCR were larger for the anticipation of completely predicted pain (100% instructions) than for unpredictable pain (25%, 50%, and 75% instructions; $p < .001$), which were larger than SCR to the anticipation of completely predicted pain omission (0% instructions, $p < .05$, except for 25% vs 0% where $p = .11$). Additionally, anticipatory responses to unpredictable pain instructions increased as a function of the provided probability instructions (75% vs 25%, $p < .05$).

A follow-up 4 (Probability: 25%, 50%, 75%, 100%) x 3 (Intensity: weak, moderate, strong) x 4 (Run: 1, 2, 3, 4) LMM examining the combined effect of Probability and Intensity in more detail showed that anticipatory SCR increased for both increasing probability ($F(3,1468) = 54.36$, $p < .001$, $\omega_p^2 = 0.10$) and increasing intensity instructions ($F(2,1468) = 67.33$, $p < .001$, $\omega_p^2 = 0.08$). Responses were overall higher to the strongest intensity ($p < .001$) compared to the weak and moderate intensities, which did not differ significantly ($p = .15$). Finally a significant Probability x Intensity interaction revealed that the effect of Probability was most pronounced when anticipating a strong stimulus with 100% certainty, and not significant for any of the 25% to 75% comparisons ($p > .07$) ($F(6,1468.06) = 2.94$, $p < .01$, $\omega_p^2 = 0.01$). Notably, the effect of Run was also significant ($F(3,1468.23) = 30.96$, $p < .001$, $\omega_p^2 = 0.06$), indicating that anticipatory SCR gradually declined over time (run 1 vs run 4, $p < .001$). Nevertheless, there were no significant interactions of Run with either Intensity or Probability ($p$'s $> .098$), suggesting that responses to the effects of interest did not alter over time.

**Supplementary Figure 4 : Individual anticipatory SCR followed probability instructions**

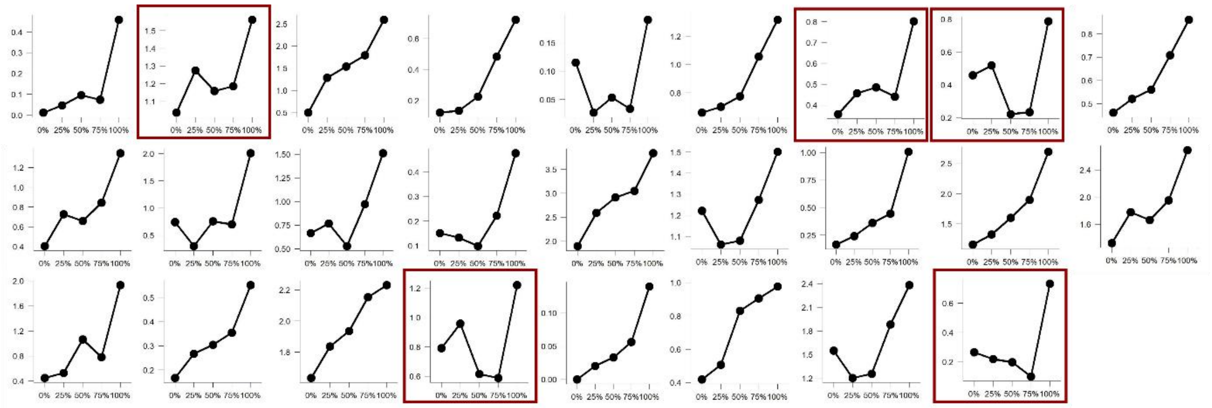


The absence of a clear increase in anticipatory SCR over 25% to 75% probability instructions on a group level, could indicate that participants did not believe the instructed probabilities. In fact, the probability instructions did not map exactly onto the actual experienced probabilities of stimulation: all instructions were followed by a stimulation in 25% of the trials. It might therefore be that the small to absent probability effects for omission responses were the result of aberrant anticipation of stimulation. In order to control for this alternative explanation, we plotted for each participant their anticipatory SCR over increasing probability levels. These plots revealed that most participants did have larger anticipatory SCR responses for 75% trials compared to 25% trials, in line with our instructions, and only 5 participants had the opposite pattern (red frames). These participants were removed in a follow-up subgroup analysis.

**Supplementary Figure 5: Run and US unpleasantness effects for subjective and physiological omission responses**

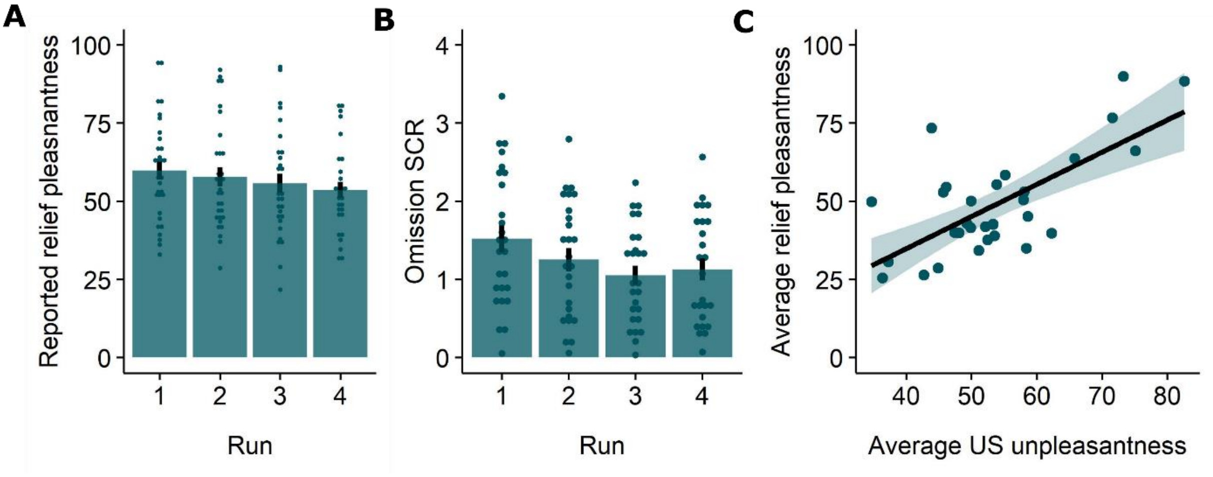


In addition to the Intensity and Probability predictors of interest, run number and average US unpleasantness (mean-centered) were entered in the LMM examining relief-pleasantness and omission SCR as predictors of no-interest. For both outcome variables we found significant main effects of Run, indicating that reported relief (**A.** $F(3,1031.73) = 9.56$, $p < .001$, $\omega_p^2 = 0.02$) and omission SCR (**B.** $F(3,862.21) = 15.51$, $p < .001$, $\omega_p^2 = 0.05$) decreased over runs (for all outcome variables, Run 1 > Run 4 contrast, $p < .001$). The overall absence of significant interactions with Probability and Intensity suggests that our effects of interest did not significantly change over time. Note, there was a trend-level Intensity x Run interaction for relief-pleasantness ($F(6,1031) = 1.98$, $p = .065$). However, post-hoc contrasts confirmed that the intensity effect did not disappear over blocks. **C.** We found that the average unpleasantness of the stimulation had a significant positive effect on the self-reported relief ($\beta = 0.96$, $p$ <.001), suggesting that the more unpleasant the stimulation was perceived, the more pleasant the relief participants reported whenever the stimulation was omitted.

**Supplementary Figure 6: Probability and effects for self-reported unpleasantness and stimulation SCR**

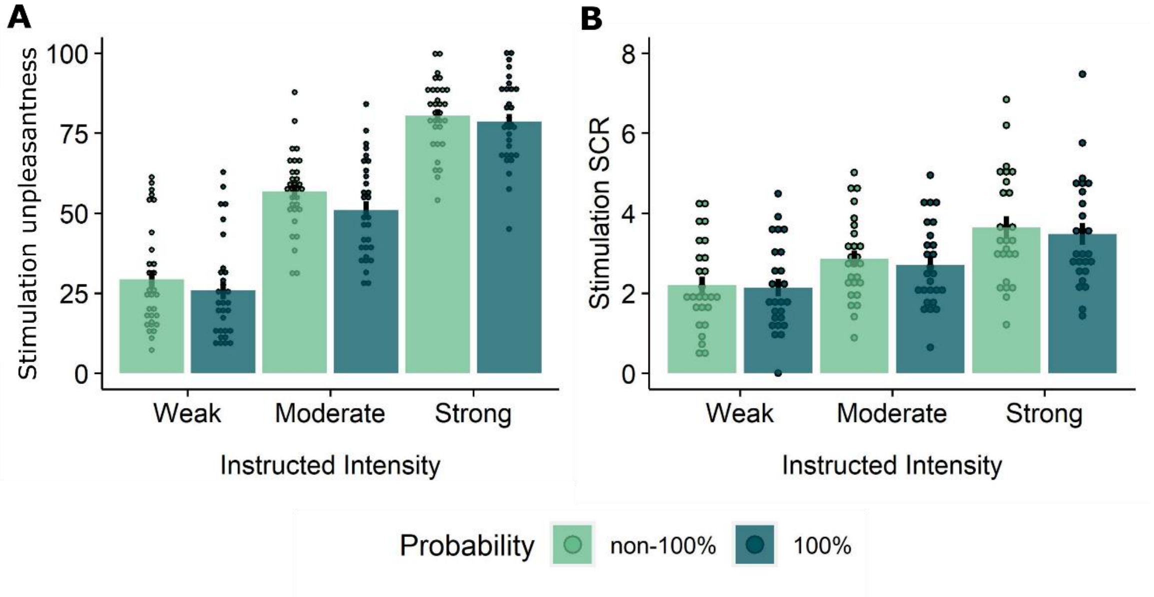


The 2 (probability: non-100%, 100%) x 3 (Intensity: weak, moderate, strong) x 4 (Run: 1, 2, 3, 4) LMM showed that unexpected stimulations (following 25%, 50%, 75% instructions) (**A)** were rated as significantly more unpleasant and (**B)** elicited higher SCR than fully expected stimulations (following 100% instructions), evidenced by main effects of Probability for self-reported unpleasantness ($F(1,678) = 16.15$, $p < .001$, $\omega_p^2 = 0.02$) and stimulation SCR ($F(1,569) = 4.07$, $p < .05$, $\omega_p^2 = 0.01$). Likewise, (**A**) stronger stimulations were experienced as more unpleasant and (**B**) elicited higher SCR, evidenced by main effects of Intensity for self-reported unpleasantness ($F(2,678) = 1015.37$, $p < .001$, $\omega_p^2 = 0.75$) and stimulation SCR ($F(2,569) = 156.33$, $p < .001$, $\omega_p^2 = 0.35$) and confirmed by follow-up Bonferroni-Holm corrected pairwise comparisons ($p$'s $< .001$).

**Supplemental Note 6: The effect of Run and the Gambler's Fallacy**

A question that was raised by the reviewers was whether omission-related responses could be influenced by dynamical learning or the Gambler's Fallacy, which might have affected the effectiveness of the Probability manipulation.

Inspired by this question, we exploratorily assessed the role of the Gambler's Fallacy and the effects of Run in a separate set of analyses. Indeed, it is possible that participants start to expect a stimulation more when more time has passed since the last stimulation was experienced. To test this alternative hypothesis, we specified two new regressors that calculated for each trial of each participant how many trials had passed since the last stimulation (or since the beginning of the experiment) either overall (across all trials of all probability types; hence called the overall-lag regressor) or per probability level (across trials of each probability type separately; hence called the lag-per-probability regressor). For both regressors a value of 0 indicates that the previous trial was either a stimulation trial or the start of experiment, a value of 1 means that the last stimulation trial was 2 trials ago, etc.

The new models including these regressors for each omission response type (i.e., omission-related activations for each ROI, relief, and omission-SCR) were specified as follows:

1. *For the overall lag*

Omission response ~ Probability * Intensity * Run + US-unpleasantness + Overall-lag + (1|Subject).

2. *For the lag per probability level*

Omission response ~ Probability * Intensity * Run + US-unpleasantness + Lag-per-probability : Probability + (1|Subject).

Where US-unpleasantness scores were mean-centered across participants; "*" represents main effects and interactions, and ":" represents an interaction (without main effect). Note that we only included an interaction for the lag-per-probability model to estimate separate lag-parameters for each probability level.

The results of these analyses are presented in the tables below. Overall, we found that adding these lag-regressors to the model did not alter our main results. That is: for the VTA/SN, relief and omission-SCR, the main effects of Probability and Intensity remained. Interestingly, the overall-lag-effect itself was significant for VTA/SN activations and omission SCR, indicating that VTA/SN activations were larger when more time had passed since the last stimulation (beta = 0.19), whereas SCR were smaller when more time had passed (beta = -0.03). This pattern is reminiscent of the Perruchet effect, namely that the explicit expectancy of a US increases over a run of non-reinforced trials (in line with the gambler's fallacy effect) whereas the conditioned physiological response to the conditional stimulus declines (in line with the extinction effect, Perruchet, 1985; McAndrew, Jones, McLaren, & McLaren, 2012). Thus, the observed dissociation between the VTA/SN activations and omission SCR might similarly point to two distinctive processes where VTA/SN activations are more dependent on a consciously controlled process that is subjected to the gambler's fallacy, whereas the strength of the SCR responses is more dependent on an automatic associative process that is subjected to extinction. Importantly, however, even though the temporal distance to the last stimulation had these opposing effects on VTA/SN activations and omission SCRs, the main effects of the probability manipulation remained significant for both outcome variables. This means that the core results of our study still hold.

Next to the overall-lag effect, the lag-per-probability regressor was only significant for the vmPFC. A follow-up of the beta estimates of the lag-per-probability regressors for each probability level revealed that vmPFC activations increased with temporal distance from shock, but only for the 50% trials (beta = 0.47 , t = 2.75, p < .01), and not the 25% (beta = 0.25, t = 1.49, p = .14) or the 75% trials (beta = 0.28, t = 1.62, p = .10).

Table 1 F-statistics and corresponding p-values from the overall lag model

| | Omission responses | | | | | | | | | | | |
|---|---|---|---|---|---|---|---|---|---|---|---|---|
| Regressor | Relief | | SCR | | VTA/SN (*) | | Left vPut | | NAC | | vmPFC | |
| | *F* | *P* | *F* | *P* | *F* | *P* | *F* | *P* | *F* | *P* | *F* | *P* |
| Probability | **30.04** | **<.001** | **4.90** | **<.01** | **3.59** | **<.05** | 0.18 | n.s. | 0.88 | n.s. | 1.73 | n.s. |
| Intensity | **620.62** | **<.001** | **106.65** | **<.001** | **7.81** | **<.001** | **3.88** | **<.05** | 0.70 | n.s. | **4.90** | **<.01** |
| Run | **9.71** | **<.001** | **15.56** | **<.001** | 1.13 | n.s. | 0.76 | n.s. | 0.62 | n.s. | 0.44 | n.s. |
| Probability x Intensity | **3.69** | **<.01** | 1.54 | n.s. | 1.15 | n.s. | 1.39 | n.s. | 1.76 | n.s. | 0.70 | n.s. |
| Probability x Run | 1.13 | n.s. | 1.24 | n.s. | 1.02 | n.s. | 1.22 | n.s. | 1.74 | n.s. | 1.26 | n.s. |
| Intensity x Run | 1.94 | .07 | 1.30 | n.s. | 1.94 | .07 | 1.59 | n.s. | 1.01 | n.s. | **2.41** | **<.05** |
| Probability x Intensity x Run | 0.56 | n.s. | 0.87 | n.s. | 0.76 | n.s. | 0.71 | n.s. | 0.76 | n.s. | 0.83 | n.s. |
| Overall-lag | 2.56 | n.s. | **4.68** | **<.05** | **11.30** | **<.001** | <0.01 | n.s. | 0.16 | n.s. | <0.01 | n.s. |
| US-unpleasantness | **29.60** | **<.001** | 3.00 | .096 | 1.84 | n.s. | 0.06 | n.s. | 3.44 | .07 | 0.26 | n.s. |

(*) F-test and p-values were based on the model where outliers were rescored to 2SD from the mean. Note that when retaining the influential outliers for this model, the p-value of the probability effect was p = .06. For all other outcome variables, rescoring the outliers did not change the results. Significant effects are indicated in bold.

Table 2 F-statistics and corresponding p-values from the lag per probability level model

| | Omission responses | | | | | | | | | | | |
|---|---|---|---|---|---|---|---|---|---|---|---|---|
| Regressor | Relief | | SCR | | VTA/SN (*) | | Left vPut | | NAC | | vmPFC | |
| | *F* | *P* | *F* | *P* | *F* | *P* | *F* | *P* | *F* | *P* | *F* | *P* |
| Probability | **23.07** | **<.001** | **4.44** | **<.05** | **3.28** | **<.05** | 0.22 | n.s. | 0.18 | n.s. | 0.31 | n.s. |
| Intensity | **625.52** | **<.001** | **107.49** | **<.001** | **7.33** | **<.001** | **3.88** | **<.05** | 0.66 | n.s. | **4.85** | **<.01** |
| Run | **8.63** | **<.001** | **13.91** | **<.001** | 1.09 | n.s. | 0.72 | n.s. | 0.60 | n.s. | 1.09 | n.s. |
| Probability x Intensity | **3.87** | **<.01** | 1.33 | n.s. | 1.01 | n.s. | 1.40 | n.s. | 1.81 | n.s. | 0.63 | n.s. |
| Probability x Run | 1.63 | n.s. | 1.10 | n.s. | 1.25 | n.s. | 1.16 | n.s. | 1.60 | n.s. | 1.17 | n.s. |
| Intensity x Run | 2.08 | .053 | 1.25 | n.s. | 1.84 | .09 | 1.61 | n.s. | 1.02 | n.s. | **2.55** | **<.05** |

| | | | | | | | | | | | | |
|---|---|---|---|---|---|---|---|---|---|---|---|---|
| Probability x Intensity x Run | 0.55 | n.s. | 0.90 | n.s. | 0.72 | n.s. | 0.71 | n.s. | 0.73 | n.s. | 0.84 | n.s. |
| Lag per probability: Probability | 1.51 | n.s. | 1.32 | n.s. | **1.12** | **n.s.** | 0.10 | n.s. | 0.96 | n.s. | **4.14** | **<.01** |
| US-unpleasantness | **29.69** | **<.001** | 2.99 | .097 | 1.63 | n.s. | 0.06 | n.s. | 3.34 | .08 | 0.27 | n.s. |

(*) F-test and p-values were based on the model where outliers were rescored to 2SD from the mean. Note that when retaining the influential outliers for this model, the p-value of the Intensity x Run interaction was p = .05. For all other outcome variables, rescoring the outliers did not change the results. Significant effects are indicated in bold.

**Supplementary fMRI results**

In addition to examining how unexpected omissions of threat are processed in the pre-specified ROIs, we explored anticipatory, omission and stimulation related whole-brain fMRI activations. Results of these analyses are presented below for exploratory purposes, but are not interpreted. For each contrast, group-level activity maps were masked with a grey matter mask and thresholded at $p < .001$ (uncorrected). An overview of the (de)activations is shown for exploratory purposes in the Supplementary Figures and MNI co-ordinates of the peak activations within each cluster can be found in the Supplementary Tables.

**Supplementary Figure 7: Whole-brain anticipatory fMRI responses to the presentation of the instructions**

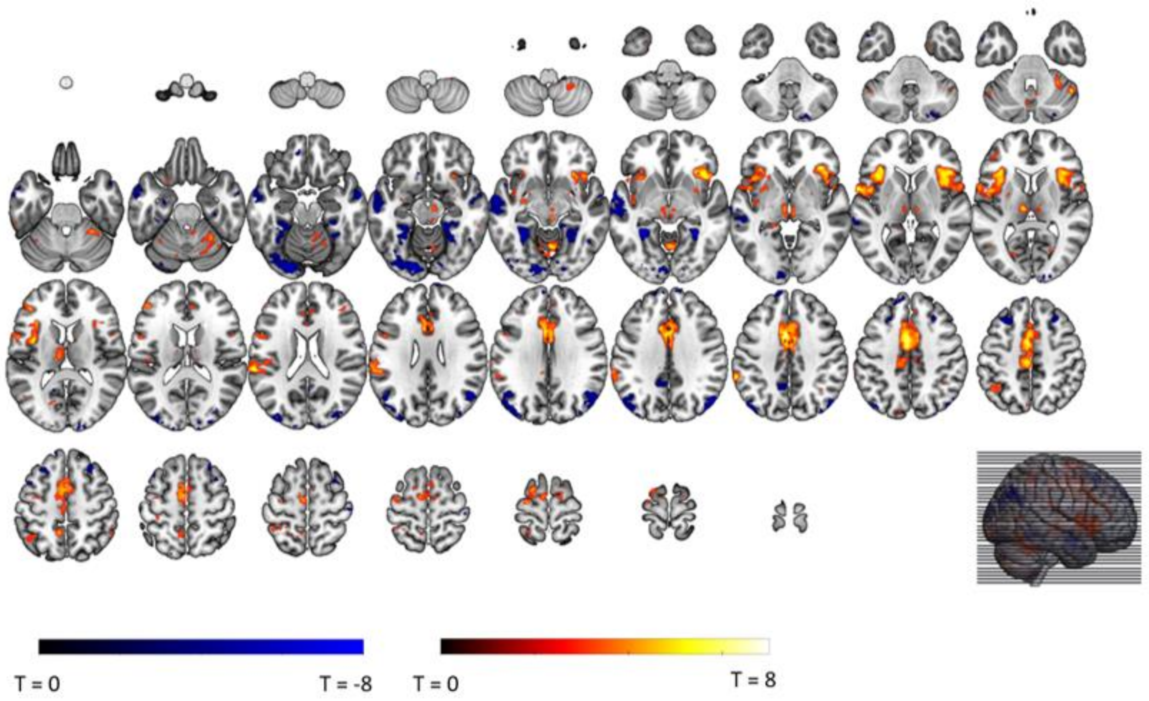


We reran our first level model, now including separate epoch-regressors for the onset of instructions of each probability x intensity combination (13 regressors), and separate general regressors for the omissions, stimulations and ratings. We contrasted all non0% trials (where there was in fact expectation of stimulation) with 0% trials in order to assess which regions show increased/decreased activation in relation to expectation of stimulation. Note that 0% trials in our task are similar to CS- trials in standard conditioning/extinction procedures, as these trials indicated that no stimulation would follow.

**Supplementary Table 3: Whole-brain anticipatory fMRI responses to the presentation of the instructions**

| **Contrast: non0% > 0%** | | | | | | | |
|---|---|---|---|---|---|---|---|
| *L/R* | *Region* | *K* | *p (cluster)* | *MNI Coordinates (xyz)* | | | *T (peak)* |
| R | Cerebellum, vermis | 149 | < .001 | 4 | -63 | -8 | 8.80 |
| L | Superior temporal gyrus | 362 | < .001 | -59 | -33 | 21 | 8.13 |
| L | Mid cingulate gyrus | 2058 | < .001 | -5 | -1 | 45 | 7.95 |
| L | Rolandic operculum | 919 | < .001 | -59 | 4 | 5 | 7.49 |
| L | Thalamus | 175 | < .001 | -5 | -21 | 1 | 7.26 |
| R | Insula | 852 | < .001 | 42 | 18 | -6 | 7.18 |
| L | Inferior frontal gyrus | 76 | .003 | -39 | 38 | 12 | 6.08 |
| R | Brain stem | 128 | < .001 | 6 | -29 | -8 | 6.03 |
| L | Insula | 50 | .033 | -41 | -13 | -4 | 5.66 |
| R | Cerebellum | 220 | < .001 | 30 | -45 | -30 | 5.56 |
| L | Precuneus | 72 | .005 | -7 | -49 | 51 | 5.36 |
| R | Supplementary motor area | 54 | .023 | 12 | -9 | 67 | 5.26 |
| L | Inferior parietal gyrus | 78 | .003 | -45 | -55 | 49 | 4.94 |
| **Contrast: non0% < 0%** | | | | | | | |
| L | Fusiform gyrus | 369 | < .001 | -31 | -45 | -6 | 8.08 |
| L | Middle temporal gyrus | 678 | < .001 | -59 | -9 | -8 | 7.80 |
| R | Angular gyrus | 329 | < .001 | 50 | -71 | 36 | 7.30 |
| R | Fusiform gyrus | 286 | < .001 | 32 | -45 | -6 | 7.24 |
| L | Fusiform gyrus | 545 | < .001 | -23 | -85 | -17 | 6.78 |
| R | Middle temporal gyrus | 109 | < .001 | 58 | -1 | -21 | 6.66 |
| L | Superior frontal gyrus | 43 | .063 | -9 | 48 | 45 | 6.14 |
| L | Angular gyrus | 398 | < .001 | -41 | -75 | 43 | 5.81 |
| R | Middle frontal gyrus | 118 | < .001 | 28 | 26 | 45 | 5.60 |
| L | Precuneus | 83 | .002 | -5 | -49 | 36 | 5.12 |
| L | Middle frontal gyrus | 154 | <.001 | -23 | 26 | 45 | 5.10 |
| R | Cerebellum | 62 | .011 | 16 | -85 | -41 | 4.35 |

*Note*. Regions are identified at voxel-level $p < .001$, and with cluster correction $p < .05$ (FWE-corrected); *L/R* indicates if the cluster (or peak of the cluster) is part of the left or right hemisphere; *Region* name is identified using the AAL atlas; *K* is the number of voxels in the cluster; *Coordinates* are the MNI coordinates of cluster peak; *T* is the value of the T-statistic of the cluster peak.

**Supplementary Figure 8: A linear increase in whole-brain anticipatory fMRI activations for increasing probability instructions**

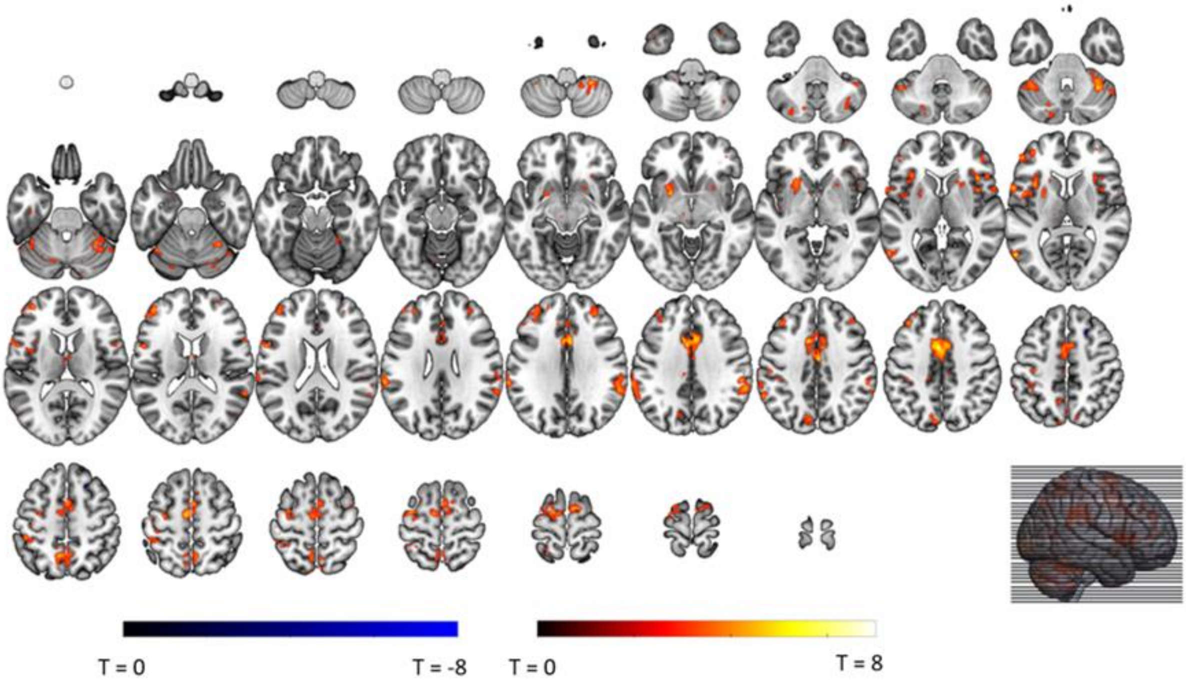


We assessed how the provided probability instructions modulated the anticipatory fMRI activations. To this end, we specified a first-level linear contrasts for probability (contrast weights: -4 (for 25%-regressors), -2 (for 50%-regressors), 2 (for 75%-regressors) and 4 (for 100% regressors)).

**Supplementary Table 4: Whole-brain anticipatory fMRI responses to increasing probability**

| **Contrast: Increase with increasing probability** | | | | | | | |
|---|---|---|---|---|---|---|---|
| *L/R* | *Region* | *K* | *p (cluster)* | *MNI Coordinates (xyz)* | | | *T (peak)* |
| L | Mid cingulate gyrus | 1394 | < .001 | -3 | 4 | 43 | 7.88 |
| L | Middle temporal gyrus | 63 | .004 | -57 | -65 | 7 | 6.74 |
| L | Precentral gyrus | 278 | < .001 | -59 | 10 | 10 | 6.65 |
| L | Putamen | 125 | < .001 | -23 | 4 | -4 | 6.30 |
| L | Postcentral gyrus | 98 | < .001 | -33 | -43 | 62 | 6.07 |
| R | Cerebellum | 64 | .004 | 28 | -45 | -50 | 6.03 |
| L | Supramarginal gyrus | 201 | < .001 | -65 | -33 | 27 | 6.02 |
| L | Middle frontal gyrus | 347 | < .001 | -41 | 46 | 16 | 5.84 |
| R | Cerebellum | 39 | .050 | 52 | -55 | -34 | 5.83 |
| R | Insula | 71 | .002 | 40 | 20 | 5 | 5.81 |
| L | Precuneus | 266 | < .001 | -9 | -55 | 54 | 5.77 |
| R | Cerebellum | 216 | < .001 | 34 | -51 | -32 | 5.72 |
| R | Middle frontal gyrus | 62 | .004 | 32 | 42 | 27 | 5.43 |
| R | Supramarginal gyrus | 277 | < .001 | 58 | -41 | 32 | 5.37 |
| L | Cerebellum | 122 | < .001 | -45 | -53 | -37 | 5.17 |
| R | Insula | 37 | .063 | 40 | 6 | 5 | 5.06 |
| L | Precuneus | 95 | < .001 | -7 | -77 | 45 | 5.06 |
| L | Cerebellum | 73 | .002 | -13 | -75 | -41 | 4.89 |
| L | Putamen | 35 | .079 | -25 | 6 | 7 | 4.65 |

| | | | | | | | |
|---|---|---|---|---|---|---|---|
| R | Cerebellum | 34 | .089 | 36 | -61 | -39 | 4.44 |
| **Contrast: Decrease with increasing probability** | | | | | | | |
| *No significant clusters of activation* | | | | | | | |

*Note.* Regions are identified at voxel-level *p* < .001, and with cluster correction *p* < .05 (FWE-corrected); *L/R* indicates if the cluster (or peak of the cluster) is part of the left or right hemisphere; *Region* name is identified using the AAL atlas; *K* is the number of voxels in the cluster; *Coordinates* are the MNI coordinates of cluster peak; *T* is the value of the T-statistic of the cluster peak.

**Supplementary Figure 9: Intensity contrast in whole-brain anticipatory fMRI activations**

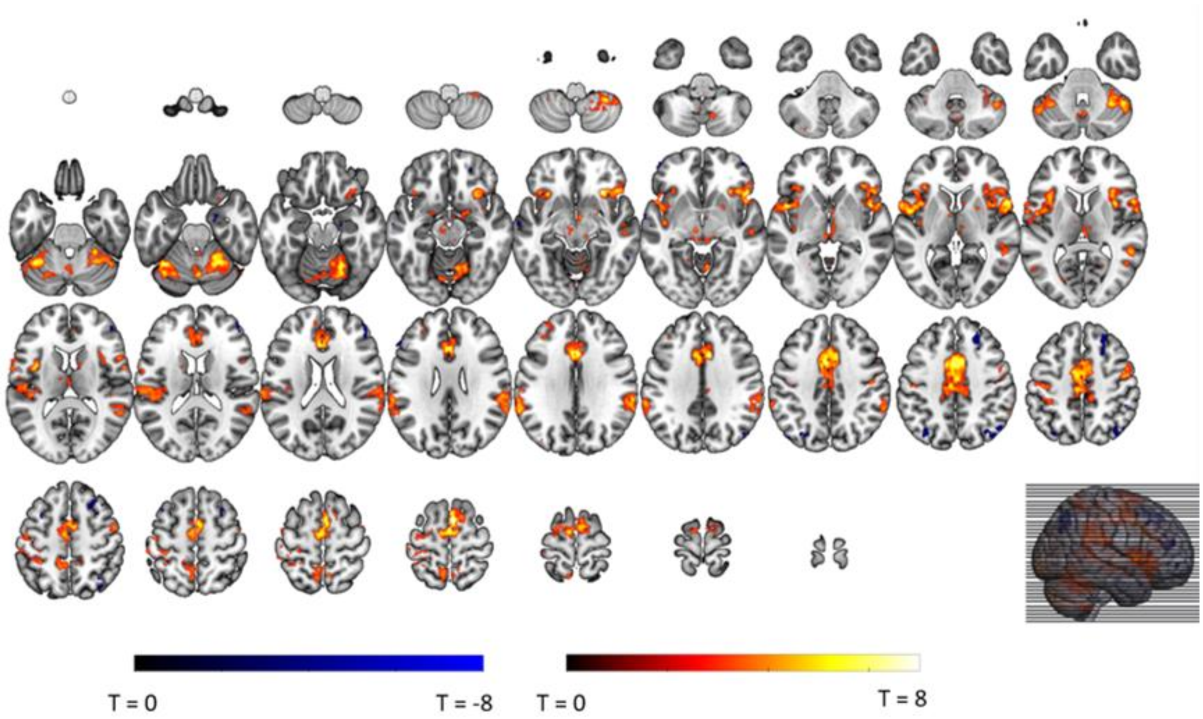


We assessed how the intensity instructions modulated the anticipatory fMRI activations. To this end, we specified a first-level contrasts for intensity (Strong > Weak).

**Supplementary Table 5: Intensity contrast in whole-brain anticipatory fMRI activations**

| **Contrast: Strong > Weak** | | | | | | | |
|---|---|---|---|---|---|---|---|
| *L/R* | *Region* | *K* | *p (cluster)* | *MNI Coordinates (xyz)* | | | *T (peak)* |
| R | Supplementary motor area | 2942 | < .001 | 6 | 6 | 67 | 9.92 |
| R | Insula | 1028 | < .001 | 30 | 22 | -8 | 8.90 |
| R | Cerebellum | 1194 | < .001 | 22 | -57 | -21 | 8.37 |
| L | Cerebellum | 478 | < .001 | -35 | -57 | -26 | 7.71 |
| L | Superior temporal gyrus | 814 | < .001 | -51 | 2 | 1 | 7.66 |
| R | Supramarginal gyrus | 674 | < .001 | 64 | -45 | 32 | 6.91 |
| L | Cerebellum | 147 | < .001 | 36 | -51 | -52 | 6.90 |
| L | Thalamus | 184 | < .001 | 6 | -17 | 1 | 6.78 |
| L | Supramarginal gyrus | 568 | < .001 | -59 | -25 | 29 | 6.48 |
| L | Postcentral gyrus | 111 | < .001 | -35 | -19 | 45 | 6.47 |
| L | Postcentral gyrus | 100 | < .001 | -31 | -43 | 62 | 6.21 |
| R | Precentral gyrus | 166 | < .001 | 52 | -7 | 47 | 6.19 |
| L | Postcentral gyrus | 75 | .002 | -31 | -31 | 54 | 5.47 |

| | | | | | | | |
|---|---|---|---|---|---|---|---|
| R | Cerebellum | 40 | .064 | 16 | -67 | -48 | 5.45 |
| L | Insula | 47 | .031 | -35 | -9 | 1 | 5.05 |
| R | Amygdala | 63 | .007 | 22 | -1 | -12 | 5.04 |
| **Contrast: Strong < Weak** | | | | | | | |
| R | Superior frontal gyrus | 170 | < .001 | 24 | 36 | 47 | 6.09 |
| R | Angular gyrus | 115 | < .001 | 40 | -73 | 45 | 5.24 |
| L | Inferior parietal gyrus | 72 | .003 | -33 | -71 | 43 | 4.80 |
| R | Middle frontal gyrus | 63 | .007 | 48 | 42 | 14 | 4.71 |

*Note*. Regions are identified at voxel-level $p < .001$, and with cluster correction $p < .05$ (FWE-corrected); *L/R* indicates if the cluster (or peak of the cluster) is part of the left or right hemisphere; *Region* name is identified using the AAL atlas; *K* is the number of voxels in the cluster; *Coordinates* are the MNI coordinates of cluster peak; *T* is the value of the T-statistic of the cluster peak.

**Supplementary Figure 10: Whole-brain omission fMRI responses**

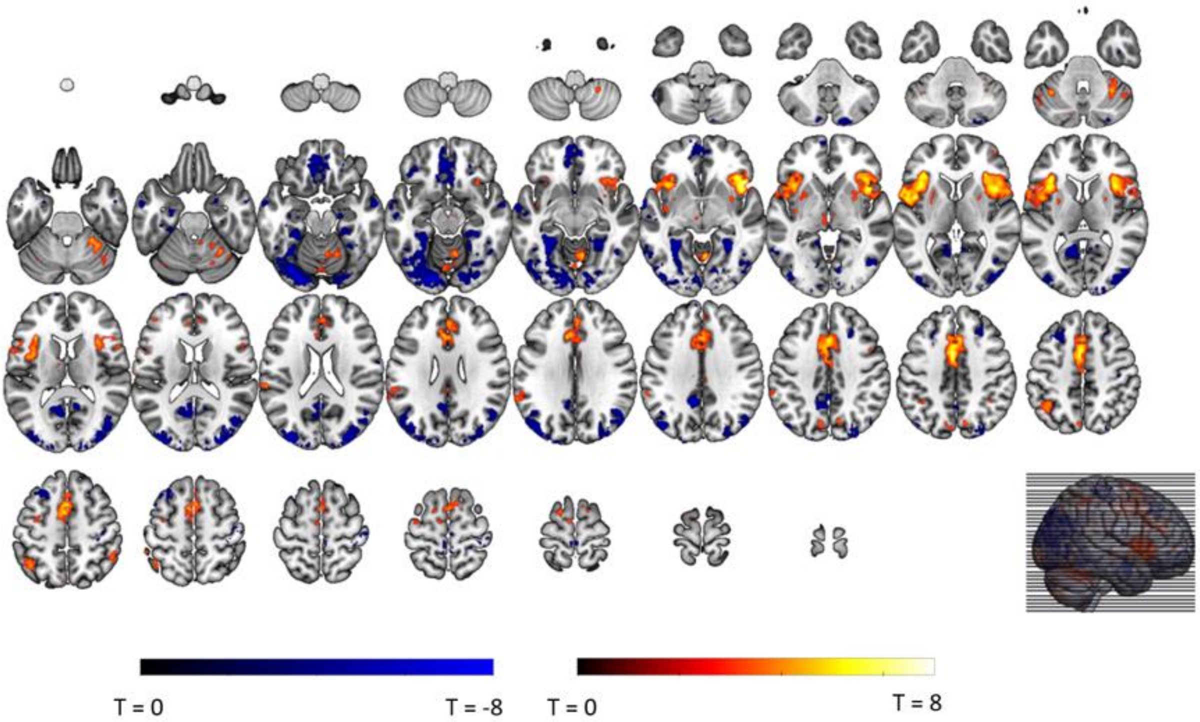


We assessed whole-brain (grey-matter masked) omission responses using the non0 % > 0% omission contrast.

**Supplementary Table 6: Whole-brain omission fMRI responses**

| L/R | Region | K | p (cluster) | MNI Coordinates (xyz) | | | T (peak) |
|---|---|---|---|---|---|---|---|
| **Contrast: non0% > 0%** | | | | | | | |
| L | Mid cingulate gyrus, extending to supplementary motor area and anterior cingulate gyrus | 1606 | < .001 | -3 | 4 | 45 | 8.19 |
| R | Insula | 1287 | < .001 | 44 | 12 | 1 | 8.05 |
| L | Insula | 1117 | < .001 | -33 | 24 | 10 | 7.45 |
| L | Putamen | 65 | .006 | -25 | 6 | 1 | 6.83 |
| R | Cerebellum | 417 | < .001 | 10 | -61 | -10 | 6.46 |
| R | Cerebellum | 37 | .093 | 38 | -63 | -26 | 5.70 |
| L | Supplementary motor area | 40 | .068 | -15 | 2 | 67 | 5.26 |
| L | Superior temporal gyrus | 148 | < .001 | -59 | -33 | 21 | 5.26 |
| L | Inferior parietal gyrus | 170 | < .001 | -43 | -53 | 54 | 5.03 |
| L | Cerebellum vermis | 55 | .016 | -3 | -75 | -15 | 4.84 |
| R | Inferior parietal gyrus | 37 | .093 | 50 | -47 | 54 | 4.74 |
| L | Supplementary motor area | 44 | .046 | -7 | -9 | 71 | 4.51 |
| **Contrast: non0% < 0%** | | | | | | | |
| L | Fusiform gyrus, extending to lingual gyrus, occipital gyrus and calcarine gyrus | 2408 | < .001 | -31 | -59 | -15 | 8.86 |
| L | Posterior cingulate gyrus | 638 | < .001 | -9 | -49 | 34 | 7.34 |
| L | vmPFC | 665 | < .001 | -5 | 42 | -17 | 6.91 |
| R | Superior occipital gyrus, extending to calcarine gyrus | 1686 | < .001 | 26 | -93 | 18 | 6.87 |
| L | Middle temporal gyrus | 176 | < .001 | -63 | -9 | -8 | 6.73 |
| R | Fusiform gyrus | 294 | < .001 | 32 | -37 | -15 | 6.73 |
| R | Precentral gyrus | 60 | .010 | 44 | -23 | 60 | 6.70 |
| R | Cerebellum | 87 | .001 | 16 | -87 | -41 | 6.33 |
| R | Middle temporal gyrus | 81 | .002 | 62 | -5 | -15 | 5.74 |
| R | Precuneus | 79 | .002 | 12 | -53 | 12 | 5.73 |
| L | Hippocampus | 64 | .007 | -25 | -11 | -23 | 5.17 |
| L | Posterior orbitofrontal gyrus | 37 | .093 | -43 | 28 | -15 | 5.12 |
| R | Middle frontal gyrus | 91 | .001 | 28 | 26 | 45 | 4.98 |
| L | Middle frontal gyrus | 230 | < .001 | -27 | 22 | 54 | 4.93 |
| L | Middle temporal gyrus | 60 | .010 | -57 | -57 | -4 | 4.89 |
| L | Paracentral lobule | 46 | .037 | -5 | -31 | 67 | 4.68 |
| R | Paracentral lobule | 39 | .076 | 6 | -37 | 62 | 4.59 |

*Note.* Regions are identified at voxel-level $p < .001$, and with cluster correction $p < .05$ (FWE-corrected); *L/R* indicates if the cluster (or peak of the cluster) is part of the left or right hemisphere; *Region* name is identified using the AAL atlas; *K* is the number of voxels in the cluster; *Coordinates* are the MNI coordinates of cluster peak; *T* is the value of the T-statistic of the cluster peak.

538 **Supplementary Figure 11: The PE-axioms within the exploratory omission clusters**

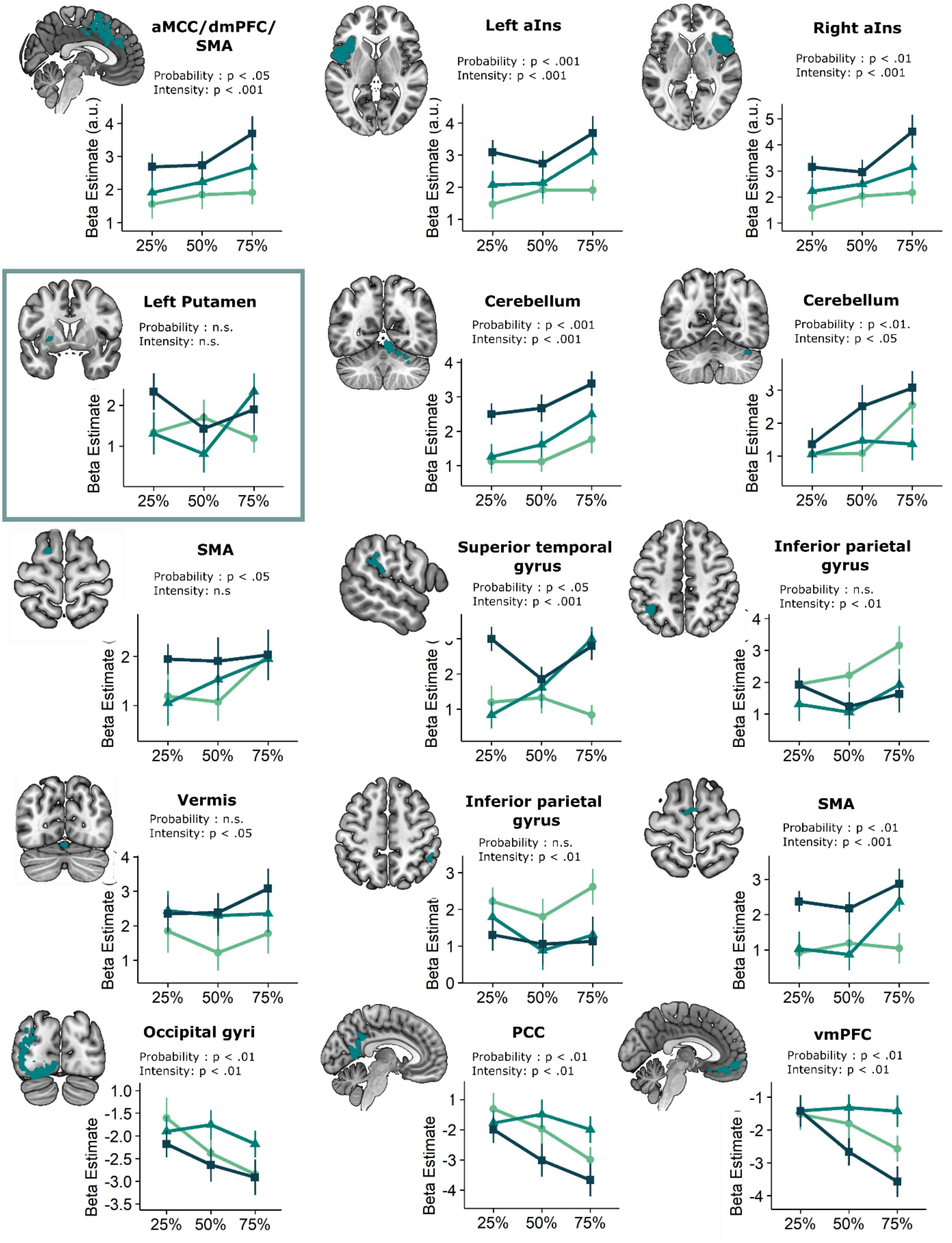


539

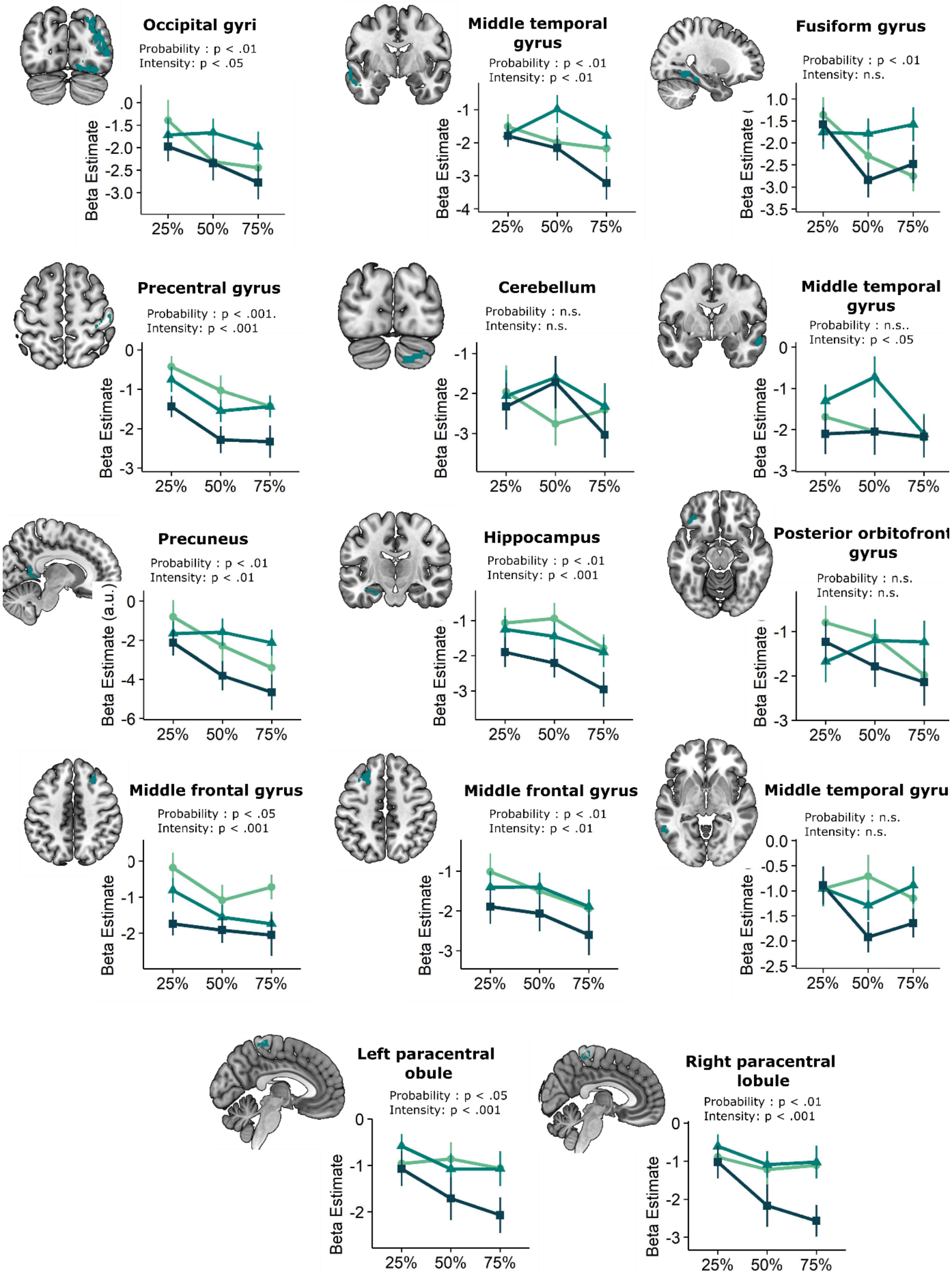


For each of the omission-processing clusters we extracted the beta-estimates per probability and intensity level. As in our main analyses, we entered them into a Probability x Intensity LMM in order to assess how omission-responses were modulated by the provided probability and intensity instructions. Results are plotted per cluster. Significance of the main effects are indicated by the reported p-values. Only the left putamen cluster did not show a difference between 100% and 0% trials (BF in favor of null-hypothesis : 3.06; axiom 3), illustrated by the teal frame. All other regions had a stronger (de)activation for 100% trials compared to 0% trials.

**Supplementary Figure 12: Whole-brain relief modulation**

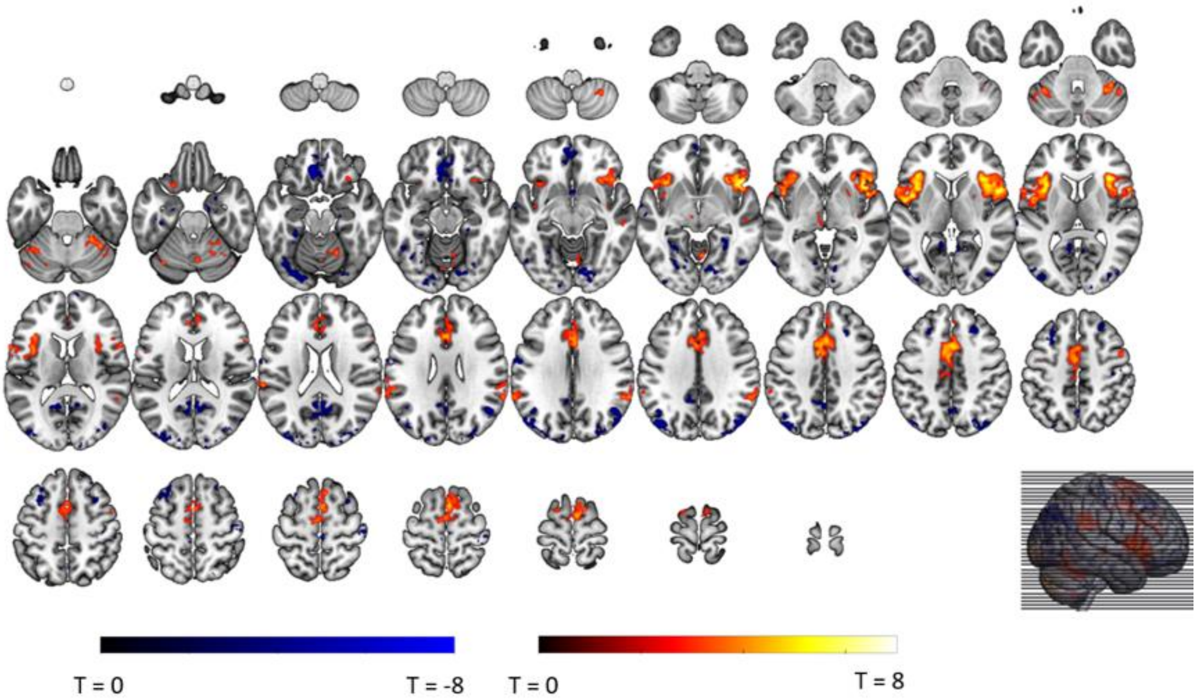


We assessed whole-brain (grey-matter masked) relief modulation of omission responses by adding trial-by-trial relief-pleasantness ratings as parametric modulator.

**Supplementary Table 7: Whole-brain relief modulation**

| L/R | Region | K | p (cluster) | MNI Coordinates (xyz) | | | T (peak) |
|---|---|---|---|---|---|---|---|
| **Contrast: Positive modulation** | | | | | | | |
| R | Insula | 1212 | < .001 | 42 | 16 | -4 | 8.91 |
| L | Insula | 1027 | < .001 | -35 | 12 | 5 | 8.31 |
| R | Mid cingulate gyrus, extending to supplementary motor area and anterior cingulate gyrus | 1739 | < .001 | 2 | 10 | 43 | 6.80 |
| R | Cerebellum | 191 | < .001 | 38 | -51 | -30 | 6.74 |
| R | Superior temporal gyrus, extending to supramarginal gyrus | 174 | < .001 | 66 | -39 | 23 | 6.71 |
| R | Cerebellum | 114 | < .001 | 8 | -71 | -12 | 6.60 |
| L | Supramarginal gyrus | 155 | < .001 | -63 | -47 | 29 | 5.84 |
| L | Cerebellum | 99 | < .001 | -37 | -51 | -32 | 5.22 |
| R | Medial frontal gyrus | 44 | .042 | 6 | 34 | 38 | 5.00 |
| **Contrast: Negative modulation** | | | | | | | |
| L | vmPFC | 381 | < .001 | -7 | 32 | -17 | 9.12 |
| R | Angular gyrus, extending to occipital gyri | 523 | < .001 | 48 | -73 | 32 | 7.59 |
| L | Angular gyrus, extending to occipital gyri | 535 | < .001 | -45 | -73 | 36 | 6.62 |
| L | Fusiform gyrus | 40 | .063 | -27 | -39 | -19 | 5.98 |
| R | Middle frontal gyrus | 107 | < .001 | 26 | 28 | 45 | 5.70 |
| R | Lingual gyrus | 167 | < .001 | 14 | -79 | -12 | 5.68 |
| R | Superior occipital gyrus | 39 | .070 | 16 | -91 | 18 | 5.40 |
| L | Precuneus | 534 | < .001 | -1 | -63 | 45 | 5.33 |
| L | Lingual gyrus | 191 | < .001 | -29 | -81 | -17 | 5.22 |
| L | Middle frontal gyrus | 194 | < .001 | -29 | 20 | 47 | 5.19 |
| L | Postcentral gyrus | 69 | .004 | -63 | -5 | 32 | 4.78 |
| L | Lingual gyrus | 48 | .028 | -27 | -53 | -6 | 4.46 |

*Note*. Regions are identified at voxel-level $p < .001$, and with cluster correction $p < .05$ (FWE-corrected); *L/R* indicates if the cluster (or peak of the cluster) is part of the left or right hemisphere; *Region* name is identified using the AAL atlas; *K* is the number of voxels in the cluster; *Coordinates* are the MNI coordinates of cluster peak; *T* is the value of the T-statistic of the cluster peak.

**Supplementary Figure 13: Whole-brain SCR modulation**

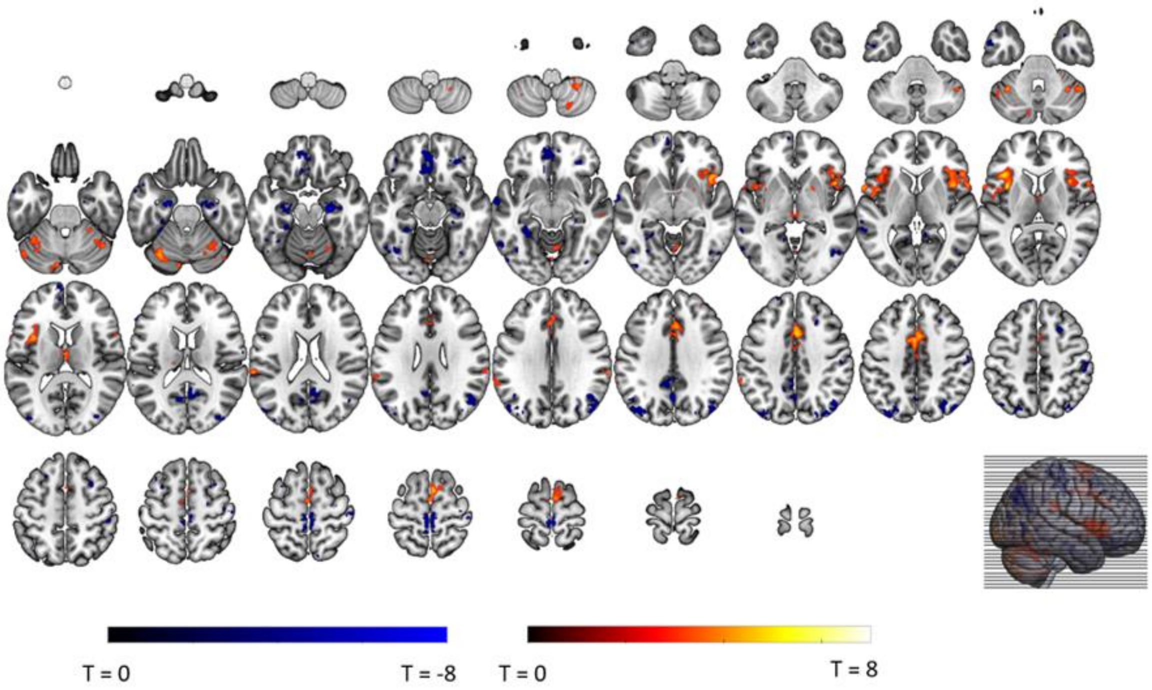


We assessed whole-brain (grey-matter masked) SCR modulation of omission responses by adding trial-by-trial SCR responses to the omission of the stimulation as parametric modulator to the omission regressor (similar to the parametric modulation analyses of relief).

**Supplementary Table 8: Whole-brain SCR modulation**

| **Contrast: Positive modulation** | | | | | | | |
|---|---|---|---|---|---|---|---|
| *L/R* | *Region* | *K* | *p (cluster)* | *MNI Coordinates (xyz)* | | | *T (peak)* |
| L | Insula | 426 | < .001 | -35 | 8 | 10 | 6.86 |
| R | Mid cingulate gyrus | 410 | < .001 | 2 | 12 | 38 | 6.03 |
| L | Superior temporal gyrus | 87 | .001 | -63 | -33 | 21 | 6.03 |
| R | Supplementary motor gyrus | 235 | < .001 | 4 | 4 | 65 | 6.01 |
| R | Insula | 554 | < .001 | 46 | 20 | -4 | 5.91 |
| L | Cerebellum | 163 | < .001 | -35 | -53 | -30 | 5.76 |
| L | Thalamus | 45 | .029 | -1 | -11 | 12 | 5.70 |
| R | Cerebellum | 50 | .017 | 30 | -49 | -50 | 5.59 |
| R | Cerebellum | 156 | < .001 | 24 | -63 | -19 | 5.56 |
| R | Cerebellum, vermis | 45 | .029 | 6 | -61 | -6 | 5.16 |
| R | Supramarginal gyrus | 35 | .087 | 64 | -33 | 29 | 5.07 |
| L | Cerebellum | 42 | .040 | -9 | -81 | -26 | 5.01 |
| L | Cerebellum | 47 | .023 | -45 | -71 | -26 | 4.90 |
| **Contrast: Negative modulation** | | | | | | | |
| L | Hippocampus | 110 | < .001 | -23 | -11 | -23 | 7.02 |
| R | Hippocampus | 153 | < .001 | 26 | -17 | -19 | 7.00 |
| L | Paracentral lobule | 123 | < .001 | -3 | -31 | 69 | 6.65 |
| R | Paracentral lobule | 128 | < .001 | 4 | -31 | 67 | 6.49 |
| L | vmPFC | 317 | < .001 | -3 | 42 | -15 | 6.38 |
| R | Angular gyrus | 332 | < .001 | 44 | -59 | 29 | 6.31 |

| | | | | | | | |
|---|---|---|---|---|---|---|---|
| L | Precuneus | 64 | .004 | -1 | -61 | 36 | 5.96 |
| L | Precuneus | 37 | .070 | -9 | -63 | 16 | 5.74 |
| R | Middle frontal gyrus | 50 | .017 | 30 | 36 | -12 | 5.67 |
| R | Posterior cingulate gyrus | 102 | < .001 | 10 | -57 | 21 | 5.61 |
| L | Fusiform gyrus | 146 | < .001 | -29 | -41 | -21 | 5.56 |
| R | Precentral gyrus | 109 | < .001 | 46 | -21 | 60 | 5.54 |
| L | Middle occipital gyrus | 274 | < .001 | -37 | -75 | 34 | 5.36 |
| R | Inferior temporal gyrus | 71 | .002 | 42 | -67 | -4 | 5.17 |
| L | Superior temporal gyrus | 52 | .014 | -63 | -5 | -6 | 5.11 |
| L | Middle temporal gyrus | 40 | .050 | -59 | -63 | -1 | 5.02 |
| L | Middle temporal gyrus | 39 | .056 | -61 | -41 | 3 | 4.91 |
| L | Middle cingulate gyrus | 89 | < .001 | -5 | -41 | 36 | 4.88 |
| R | Middle frontal gyrus | 58 | .008 | 26 | 18 | 47 | 4.79 |
| L | Inferior temporal gyrus | 56 | .009 | -47 | 2 | -41 | 4.54 |

*Note*. Regions are identified at voxel-level $p < .001$, and with cluster correction $p < .05$ (FWE-corrected); *L/R* indicates if the cluster (or peak of the cluster) is part of the left or right hemisphere; *Region* name is identified using the AAL atlas; *K* is the number of voxels in the cluster; *Coordinates* are the MNI coordinates of cluster peak; *T* is the value of the T-statistic of the cluster peak.

**Supplementary Figure 14 : The neural signature of omission SCR – a LASSO-PCR analysis of omission SCR**

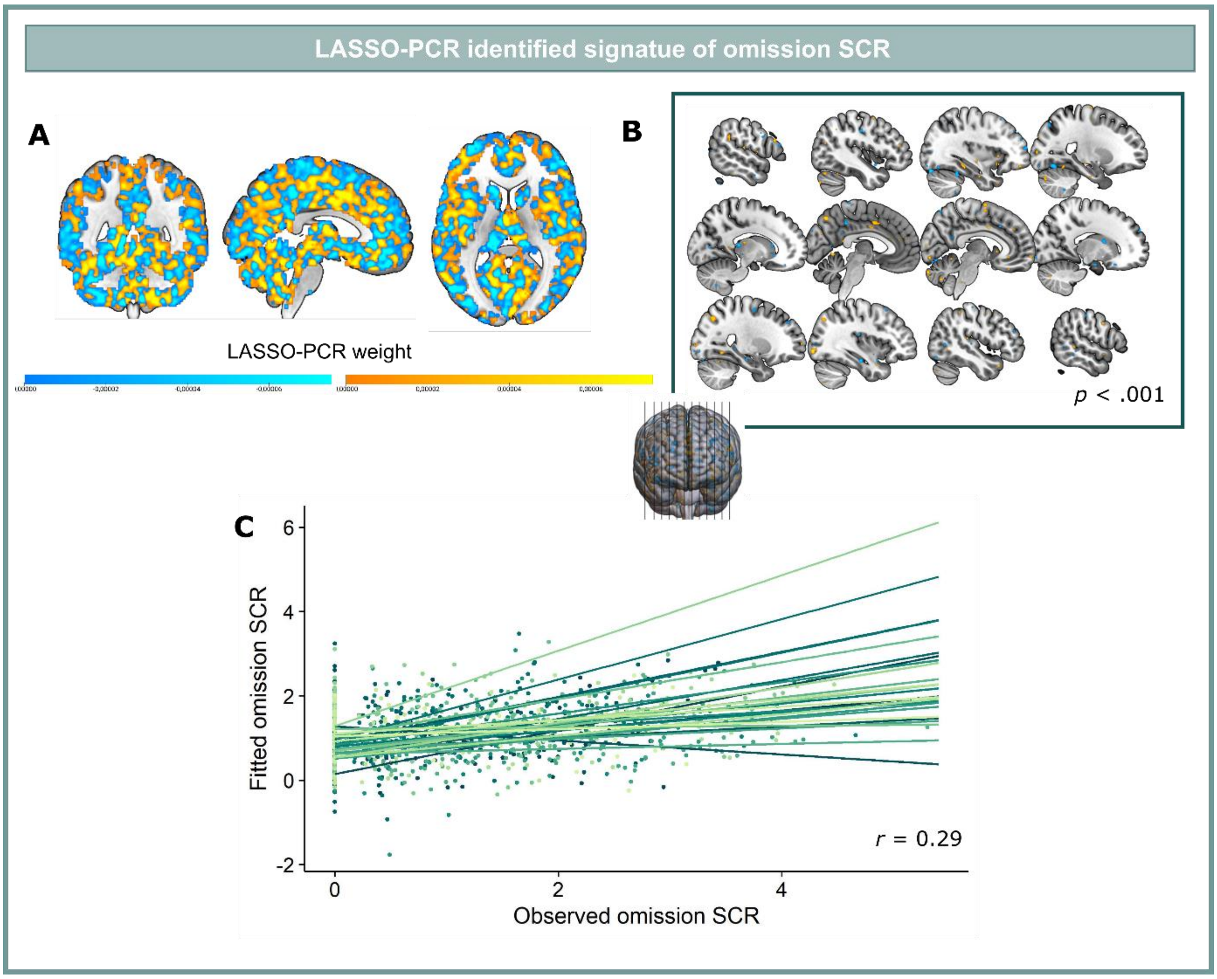

Similar to the LASSO-PCR model of relief, we trained a LASSO-PCR model for omission SCR in order to identify the pattern of omission-related brain responses that can predict the magnitude of the omission SCR response. The model was trained based on the data of SCR-responders (N = 25), using five-fold cross validation. **(A)** This yielded a map of positive and negative regression weights (signature response). **(C)** Predicted and observed SCR correlated significantly ($r$ = .29, $p$ < .001). **(B)** Bootstrap tests (5000 samples) were identify the features that contributed most the prediction.

**Supplementary Table 9: Main omission SCR signature clusters identified via bootstrapping**

| **Positive weight clusters** | | | | | | |
|---|---|---|---|---|---|---|
| *L/R* | *Region* | *K* | *MNI Coordinates (xyz)* | | | *Z (peak)* |
| R | Cerebellum Crus 2 | 11 | 8 | -87 | -30 | 0.00019 |
| L | Cerebellum Crus 2 | 12 | -19 | -87 | -28 | 0.00016 |
| L | Cerebellum Crus 1 | 16 | -41 | -71 | -26 | 0.00015 |
| R | Calcarine gyrus | 12 | 2 | -91 | -12 | 0.00023 |
| R | Fusiform gyrus | 10 | 30 | -57 | -8 | 0.00020 |
| R | Inferior occipital gyrus | 15 | 36 | -89 | -6 | 0.00023 |
| L | Anterior cingulate cortex | 12 | -1 | 26 | 18 | 0.00026 |
| L | Supramarginal gyrus | 21 | -55 | -45 | 27 | 0.00020 |
| L | Cuneus | 12 | -9 | -81 | 29 | 0.00018 |
| R | Superior occipital gyrus | 70 | 30 | -71 | 43 | 0.00018 |
| L | Postcentral gyrus | 12 | -61 | -19 | 36 | 0.00013 |
| L | Precuneus | 21 | -3 | -71 | 51 | 0.00023 |
| **Negative weight clusters** | | | | | | |
| L | Cerebellum | 18 | -33 | -45 | -26 | -0.00025 |
| L | Cerebellum Crus 1 | 11 | -35 | -83 | -21 | -0.00031 |
| R | Lingual gyrus | 12 | 30 | -85 | -17 | -0.00026 |
| L | Fusiform gyrus | 33 | -23 | -69 | -15 | -0.00025 |
| R | Superior frontal gyrus | 16 | 40 | 42 | -15 | -0.00020 |
| L | Lingual gyrus | 15 | -39 | -85 | -12 | -0.00014 |
| R | Inferior occipital gyrus | 29 | 42 | -79 | -12 | -0.00014 |
| R | Inferior temporal gyrus | 10 | 60 | -41 | -12 | -0.00015 |
| R | Inferior occipital gyrus | 13 | 30 | -89 | -6 | -0.00015 |
| R | Middle temporal gyrus | 11 | 52 | -55 | 1 | -0.00017 |
| L | Middle occipital gyrus | 24 | -39 | -87 | 5 | -0.00016 |
| R | Middle occipital gyrus | 17 | 40 | -85 | 5 | -0.00019 |
| R | Middle frontal gyrus | 17 | 42 | 42 | 7 | -0.00014 |
| L | Middle occipital gyrus | 17 | -51 | -73 | 7 | -0.00020 |
| L | Caudate | 11 | -17 | 20 | 7 | -0.00019 |
| L | Thalamus | 16 | -13 | -31 | 10 | -0.00026 |
| R | Calcarine gyrus | 10 | 4 | -75 | 12 | -0.00019 |
| R | Caudate | 14 | 14 | 6 | 16 | -0.00016 |
| R | Postcentral gyrus | 10 | 62 | -3 | 21 | -0.00017 |
| R | Frontal middle gyrus | 17 | 44 | 40 | 23 | -0.00015 |
| R | Superior frontal gyrus | 11 | 20 | 62 | 23 | -0.00015 |
| L | Precentral gyrus | 11 | -55 | 6 | 29 | -0.00013 |
| R | Precuneus | 10 | 2 | -77 | 45 | -0.00019 |
| L | Postcentral gyrus | 11 | -51 | -15 | 54 | -0.00016 |
| R | Precuneus | 11 | 10 | -69 | 60 | -0.00021 |
| R | Supplementary motor area | 15 | 4 | -25 | 60 | -0.00015 |

*Note*. Clusters (FDR-corrected, k >10, following bootstrapping). *L/R* indicates if the cluster (or peak of the cluster) is part of the left or right hemisphere; *Region* name is identified using the AAL atlas. *K* is the number of voxels in the cluster; *coordinates* are the MNI coordinates of cluster peak, Z is the signature weight of the cluster peak.

**Supplementary Figure 15: Stimulation fMRI results**

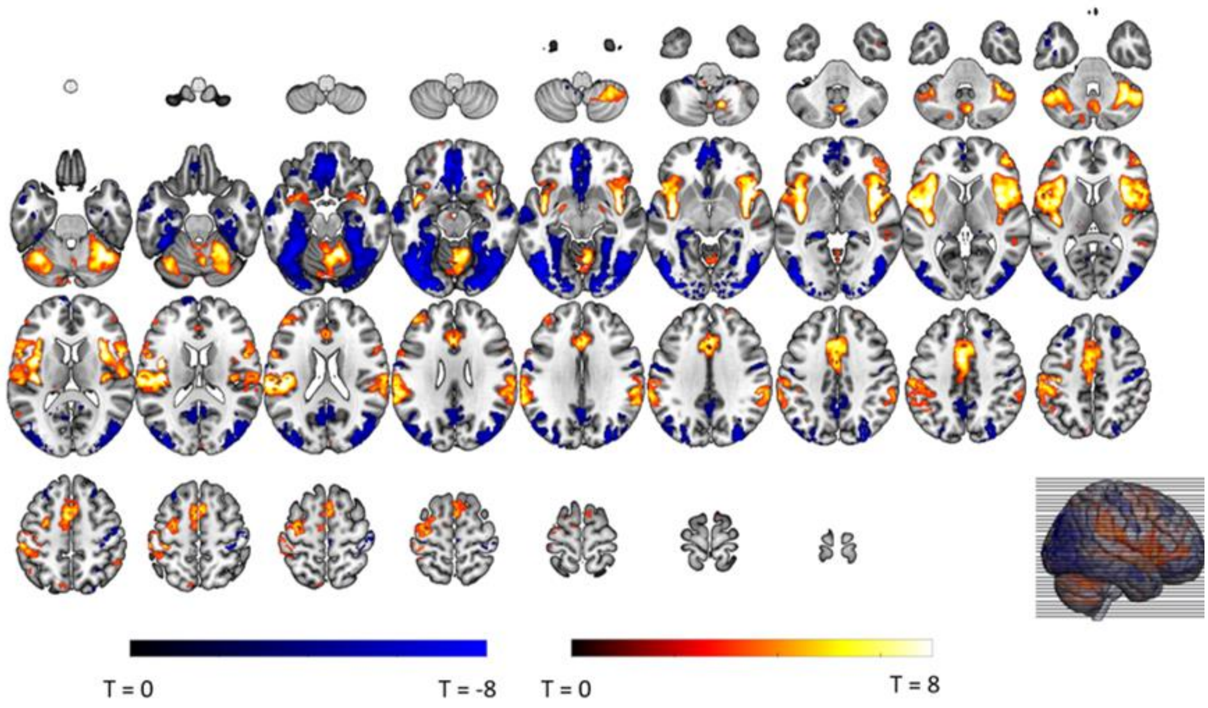


We assessed whole-brain (grey-matter masked) stimulation-related fMRI responses based on the stimulation > baseline contrast.

**Supplementary Table 10: Whole-brain stimulation-induced activations**

| **Contrast: Stimulation > baseline** | | | | | | | |
|---|---|---|---|---|---|---|---|
| *L/R* | *Region* | *K* | *p (cluster)* | *MNI Coordinates (xyz)* | | | *T (peak)* |
| L | Insula | 5078 | < .001 | -39 | -1 | 16 | 15.62 |
| R | Insula | 3774 | < .001 | 44 | 10 | -4 | 11.71 |
| R | Cerebellum | 2577 | < .001 | 14 | -55 | -17 | 11.32 |
| L | Cerebellum | 684 | < .001 | -35 | -63 | -26 | 9.92 |
| L | Mid cingulate gyrus | 2048 | < .001 | -3 | 8 | 43 | 9.44 |
| L | Middle frontal gyrus | 271 | < .001 | -41 | 42 | 23 | 7.12 |
| R | Middle temporal gyrus | 119 | < .001 | 54 | -31 | 1 | 5.11 |
| L | Cerebellum | 78 | .007 | -17 | -77 | -37 | 4.85 |
| **Contrast: Stimulation < baseline** | | | | | | | |
| R | Occipital gyri, extending to fusiform, lingual gyri | 4348 | < .001 | 32 | -91 | -10 | 14.53 |
| L | Occipital gyri, extending to fusiform, lingual gyri | 4411 | < .001 | -35 | -91 | -15 | 13.70 |
| L | vmPFC | 1883 | < .001 | -5 | 42 | -15 | 11.66 |
| R | Precentral gyrus | 271 | < .001 | 40 | -21 | 51 | 10.21 |
| R | Postcentral gyrus | 85 | .004 | 36 | -31 | 56 | 8.25 |
| R | Middle temporal gyrus | 188 | < .001 | 60 | -3 | -19 | 7.95 |

| | | | | | | | |
|---|---|---|---|---|---|---|---|
| L | Middle temporal gyrus | 582 | < .001 | -61 | -13 | -15 | 6.69 |
| R | Superior frontal gyrus | 136 | < .001 | 26 | 32 | 49 | 6.22 |
| L | Precentral gyrus | 143 | < .001 | -57 | -3 | 29 | 6.20 |
| L | Posterior cingulate gyrus | 887 | < .001 | -3 | -55 | 29 | 6.11 |
| L | Superior frontal gyrus | 136 | < .001 | -27 | 26 | 56 | 5.94 |
| R | Cerebellum | 46 | .083 | 20 | -83 | -41 | 5.31 |

*Note*. Regions are identified at voxel-level $p < .001$, and with cluster correction $p < .05$ (FWE-corrected); *L/R* indicates if the cluster (or peak of the cluster) is part of the left or right hemisphere; *Region* name is identified using the AAL atlas; *K* is the number of voxels in the cluster; *Coordinates* are the MNI coordinates of cluster peak; *T* is the value of the T-statistic of the cluster peak.

**Supplementary Figure 16: Unexpected stimulation fMRI results**

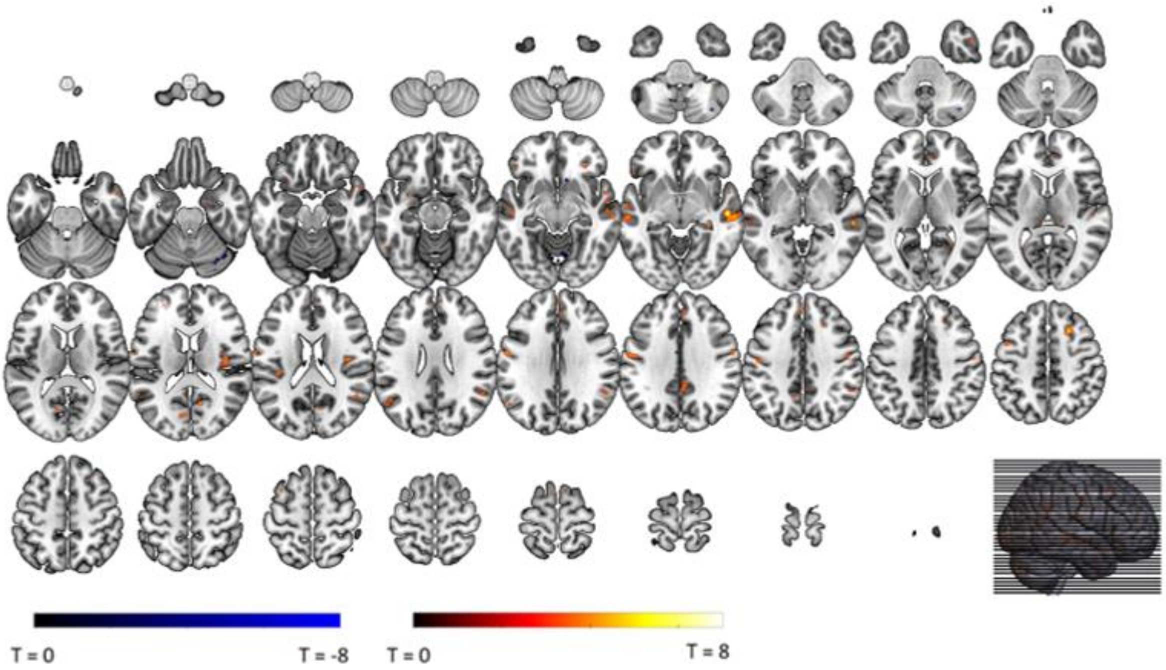


We assessed whole-brain (grey-matter masked) unexpected stimulation-related fMRI responses based on the non-100% stimulation > 100% stimulation contrast.

**Supplementary Table 11: Whole-brain unexpected stimulation-induced activations**

| **Contrast: non100% > 100% stimulations** | | | | | | | |
|---|---|---|---|---|---|---|---|
| *L/R* | *Region* | *K* | *p (cluster)* | *MNI Coordinates (xyz)* | | | *T (peak)* |
| R | Middle temporal gyrus | 121 | <.001 | 54 | -21 | -6 | 6.52 |
| R | Rolandic operculum | 50 | .014 | 42 | -11 | 18 | 5.57 |
| R | Frontal superior gyrus | 44 | .027 | 24 | 20 | 47 | 5.17 |
| L | Middle temporal gyrus | 42 | .034 | -59 | -25 | -4 | 5.10 |
| L | Precentral gyrus | 72 | .002 | -57 | -7 | 32 | 4.75 |
| R | Middle temporal gyrus | 36 | .067 | 50 | -55 | 21 | 4.67 |
| R | Medial superior frontal gyrus | 34 | .085 | 8 | 54 | 23 | 4.49 |
| R | Mid cingulate gyrus | 34 | .085 | 4 | -47 | 34 | 4.32 |
| L | Middle temporal gyrus | 33 | .096 | -53 | -55 | 23 | 4.32 |
| **Contrast: non100% < 100% stimulations** | | | | | | | |

| |
|---|
| *No significant clusters of activation* |

*Note*. Regions are identified at voxel-level $p < .001$, and with cluster correction $p < .05$ (FWE-corrected); *L/R* indicates if the cluster (or peak of the cluster) is part of the left or right hemisphere; *Region* name is identified using the AAL atlas; *K* is the number of voxels in the cluster; *Coordinates* are the MNI coordinates of cluster peak; *T* is the value of the T-statistic of the cluster peak.

Gorgolewski, K., C. D. Burns, C. Madison, D. Clark, Y. O. Halchenko, M. L. Waskom, and S. Ghosh.

2011. “Nipype: A Flexible, Lightweight and Extensible Neuroimaging Data Processing Framework in Python.” *Frontiers in Neuroinformatics* 5: 13. https://doi.org/10.3389/fninf.2011.00013.

Gorgolewski, Krzysztof J., Oscar Esteban, Christopher J. Markiewicz, Erik Ziegler, David Gage Ellis, Michael Philipp Notter, Dorota Jarecka, et al. 2018. “Nipype.” *Software*. Zenodo. https://doi.org/10.5281/zenodo.596855.

Greve, Douglas N, and Bruce Fischl. 2009. “Accurate and Robust Brain Image Alignment Using Boundary-Based Registration.” *NeuroImage* 48 (1): 63-72. https://doi.org/10.1016/j.neuroimage.2009.06.060.

Hendriks, M. H. A., Daniels, N., Pegado, F., & de Beeck, H. P. O. (2017). The effect of spatial smoothing on representational similarity in a simple motor paradigm. *Frontiers in Neurology*, *8*(MAY), 1–11. https://doi.org/10.3389/fneur.2017.00222

Hikosaka, O. (2010). The habenula: From stress evasion to value-based decision-making. *Nature Reviews Neuroscience*, *11*(7), 503–513. https://doi.org/10.1038/nrn2866

Jenkinson, Mark, Peter Bannister, Michael Brady, and Stephen Smith. 2002. “Improved Optimization for the Robust and Accurate Linear Registration and Motion Correction of Brain Images.” *NeuroImage* 17 (2): 825–41. https://doi.org/10.1006/nimg.2002.1132.

Jenkinson, Mark, and Stephen Smith. 2001. “A Global Optimisation Method for Robust Affine Registration of Brain Images.” *Medical Image Analysis* 5 (2): 143–56. https://doi.org/10.1016/S1361-8415(01)00036-6.

Klooster, P. M. ten., Weekers, A. M., Eggelmeijer, F., van Woerkom, J. M., Drossaert, C. H. C. M., Taal, E., Rasker, J. J., & Baneke, J. J. (2010). Optimisme en/of pessimisme: factorstructuur van de Nederlandse Life Orientation Test-Revised. *Psychologie En Gezondheid*, *38*(2), 89–100. https://doi.org/10.1007/BF03089356

Lanczos, C. 1964. “Evaluation of Noisy Data.” *Journal of the Society for Industrial and Applied Mathematics Series B Numerical Analysis* 1 (1): 76–85. https://doi.org/10.1137/0701007.

Linnman, C., Rougemont-Bücking, A., Beucke, J. C., Zeffiro, T. A., & Milad, M. R. (2011). Unconditioned responses and functional fear networks in human classical conditioning. *Behavioural Brain Research*, *221*(1), 237–245. https://doi.org/10.1016/j.bbr.2011.02.045

Lovibond, P. F., & Lovibond, S. H. (1995). The structure of negative emotional states: Comparison of the Depression Anxiety Stress Scales (DASS) with the Beck Depression and Anxiety Inventories. *Behaviour Research and Therapy*, *33*(3), 335–343. https://doi.org/https://doi.org/10.1016/0005-7967(94)00075-U

Op de Beeck, H. P. (2010). Against hyperacuity in brain reading: Spatial smoothing does not hurt multivariate fMRI analyses? *NeuroImage*, *49*(3), 1943–1948. https://doi.org/10.1016/j.neuroimage.2009.02.047

Power, Jonathan D., Anish Mitra, Timothy O. Laumann, Abraham Z. Snyder, Bradley L. Schlaggar, and Steven E. Petersen. 2014. “Methods to Detect, Characterize, and Remove Motion Artifact in Resting State fMRI.” *NeuroImage* 84 (Supplement C): 320–41. https://doi.org/10.1016/j.neuroimage.2013.08.048.

Satterthwaite, Theodore D., Mark A. Elliott, Raphael T. Gerraty, Kosha Ruparel, James Loughead, Monica E. Calkins, Simon B. Eickhoff, et al. 2013. “An improved framework for confound regression and filtering for control of motion artifact in the preprocessing of resting-state functional connectivity data.” *NeuroImage* 64 (1): 240–56. https://doi.org/10.1016/j.neuroimage.2012.08.052.

Scheier, M. F., Carver, C. S., & Bridges, M. W. (1994). Distinguishing Optimism From Neuroticism (and Trait Anxiety, Self-Mastery, and Self-Esteem): A Reevaluation of the Life Orientation Test.

*Journal of Personality and Social Psychology*, *67*(6), 1063–1078. https://doi.org/10.1037/0022-3514.67.6.1063

Simons, J. S., & Gaher, R. M. (2005). The distress tolerance scale: Development and validation of a self-report measure. *Motivation and Emotion*, *29*(2), 83–102. https://doi.org/10.1007/s11031-005-7955-3

Spielberger, C., Gorsuch, R., Lushene, R., Vagg, P. R., & Jacobs, G. (1983). Manual for the State-Trait Anxiety Inventory (Form Y1 – Y2). In *Palo Alto, CA: Consulting Psychologists Press; Vol. IV*.

Sullivan, M. J. L., Bishop, S. R., & Pivik, J. (1995). The Pain Catastrophizing Scale: Development and validation. *Psychological Assessment*, *7*(4), 524–532. https://doi.org/https://doi.org/10.1037/1040-3590.7.4.524

Taylor, S., Zvolensky, M. J., Cox, B. J., Deacon, B., Heimberg, R. G., Ledley, D. R., Abramowitz, J. S., Holaway, R. M., Sandin, B., Stewart, S. H., Coles, M., Eng, W., Daly, E. S., Arrindell, W. A., Bouvard, M., & Cardenas, S. J. (2007). Robust Dimensions of Anxiety Sensitivity: Development and Initial Validation of the Anxiety Sensitivity Index-3. *Psychological Assessment*, *19*(2), 176–188. https://doi.org/10.1037/1040-3590.19.2.176

Tustison, N. J., B. B. Avants, P. A. Cook, Y. Zheng, A. Egan, P. A. Yushkevich, and J. C. Gee. 2010. "N4ITK: Improved N3 Bias Correction." *IEEE Transactions on Medical Imaging* 29 (6): 1310–20. https://doi.org/10.1109/TMI.2010.2046908.

Van der Ploeg, H. M. (1982). De zelf-beoordelings vragenlijst (STAY-DY). *Tijdschrift Voor Psychiatrie*, *24*, 576–588.

Watson, D., Clark, L. A., & Tellegen, A. (1988). Development and validation of brief measures of positive and negative affect: The PANAS scales. *Journal of Personality and Social Psychology*, *54*(6), 1063–1070.

White, T. L., & Carver, C. S. (1994). Behavioral inhibition, behavioral activation, and affective responses to impending reward and punishment: The BIS/BAS scales. *Journal of Personality and Social Psychology*, *67*(2), 319–333. http://www.psy.miami.edu/faculty/ccarver/sclBISBAS.html%5Cnhttp://www.scribd.com/doc/8760706/Carver1994?secret_password=zkhvhvj8dknjmpqjutf

Zhang, Y., M. Brady, and S. Smith. 2001. "Segmentation of Brain MR Images Through a Hidden Markov Random Field Model and the Expectation-Maximization Algorithm." *IEEE Transactions on Medical Imaging* 20 (1): 45–57. https://doi.org/10.1109/42.906424.
